# The Mechanosensitive Ion Channel MSL10 Potentiates Responses to Cell Swelling in Arabidopsis Seedlings

**DOI:** 10.1101/2020.04.02.021683

**Authors:** Debarati Basu, Elizabeth S. Haswell

## Abstract

The ability to respond to unanticipated increases in volume is a fundamental property of cells, essential for cellular integrity in the face of osmotic challenges. Plants must manage cell swelling during flooding, rehydration, and pathogenesis–but little is known about the mechanisms by which this occurs. It has been proposed that plant cells could sense and respond to cell swelling through the action of mechanosensitive ion channels. Here we develop and characterize a new assay to study the effects of cell swelling on *Arabidopsis thaliana* seedlings and to test the contributions of the mechanosensitive ion channel MscS-Like10 (MSL10). The assay incorporates both cell wall softening and hypo-osmotic treatment to induce cell swelling. We show that MSL10 is required for previously demonstrated responses to hypo-osmotic shock, including a cytoplasmic calcium transient within the first few seconds, accumulation of ROS within the first 30 minutes, and increased transcript levels of mechano-inducible genes within 60 minutes. We also show that cell swelling induces programmed cell death within 3 hours, also in a MSL10-dependent manner. Finally, we show that MSL10 is unable to potentiate cell swelling-induced death when phosphomimetic residues are introduced into its soluble N-terminus. Thus, MSL10 functions as a phospho-regulated membrane-based sensor that connects the perception of cell swelling to a downstream signaling cascade and programmed cell death.

## INTRODUCTION

Cells from all organisms must actively maintain an optimal volume; left uncontrolled, the passive diffusion of water into the cell would lead to extreme swelling and loss of integrity (Wood, 2011; Hýskova and Ryslava, 2018; Toft-Bertelsen et al., 2018). Furthermore, in plants, cell volume regulation is required to maintain cell shape, accomplish cell expansion and growth, drive cellular movements like stomatal opening, and prevent wilting (Zonia and Munnik, 2007; Scharwies and Dinneny, 2019). Plant cells may swell in the early stages of flooding, rehydration after desiccation, during water soaking in pathogenesis, and in response to cell wall damage by pathogens. In the laboratory, cell swelling is induced by subjecting tissues, cell suspensions or protoplasts to sudden, large drops in media osmolarity or disrupting of microtubule cytoskeleton (for example, Sugimoto et al., 2003; Droillard et al., 2004).

There are both immediate and delayed responses to cell swelling. To immediately protect cell integrity and prevent lysis, microbes release osmolytes through channels or transporters; this raises the osmotic potential of the cell and reduces the entry of additional water (Wood, 2011; Hohmann, 2015; Cox et al., 2018). Mammalian and plant cells respond by increasing membrane area by adjusting the relative rates of endocytosis/exocytosis to favor an increase in membrane volume, thereby preventing lysis (Diz-Muñoz et al., 2013; Zwiewka et al., 2015). These rapid responses are only the beginning of the plant cell’s response to cell swelling. A transient increase in cytoplasmic calcium ([Ca^2+^]_cyt_) in response to cell swelling has been documented in cultured tobacco, rice and Arabidopsis cells (Takahashi et al., 1997b; Cazalé et al., 1998; Cessna et al., 1998; Cessna and Low, 2001; Pauly et al., 2001; Nakagawa et al., 2007; Kurusu et al., 2012a; Kurusu et al., 2012b; Nguyen et al., 2018), Arabidopsis leaves (Hayashi et al., 2006), and Arabidopsis root tips (Shih et al., 2014). Cell swelling also produces a burst of reactive oxygen species (ROS) (Yahraus et al., 1995; Beffagna et al., 2005; Rouet et al., 2006) and leads to the activation of mitogen-activated kinases (Takahashi et al., 1997a; Cazalé et al., 1998; Droillard et al., 2004). Many other stress responses to hypo-osmotic shock have also been documented (Felix et al., 2000; Liu et al., 2001; Ludwig et al., 2005; Tsugama et al., 2012). In Arabidopsis, a number of mechano- and Ca^2+-^responsive genes are induced in root tips after hypo-osmotic treatment (Shih et al., 2014). A family of bZIP transcription factors are translocated to the nucleus upon hypo-osmotic shock, where they are thought to up-regulate rehydration-responsive genes, including several enzymes that reduce levels of the plant hormone ABA (Tsugama et al., 2012; Tsugama et al., 2014; Tsugama et al., 2016). Despite these many observations, it is still not clear how these downstream signaling components are connected, nor how they might lead to adaptive responses.

It is also poorly understood how cell swelling is initially perceived in plants. The many potential signals include molecular crowding, membrane tension, osmotic gradient, turgor, membrane curvature, cell wall damage, and the disruption of the plasma membrane-cell wall connections; (Haswell and Verslues, 2015; Cuevas-Velazquez and Dinneny, 2018; Hýskova and Ryslava, 2018; Gigli-Bisceglia et al., 2019; Le Roux et al., 2019). Several families of receptor-like kinases, including the WALL-ASSOCATED KINASES, *Catharanthus roseus* RLK family, have been implicated in sensing cell wall integrity (Nissen et al., 2016; Gigli-Bisceglia et al., 2019). In particular, the *Cr*RLK FERONIA has been implicated in the perception of mechanical signals, including hypo-osmotic shock ((Shih et al., 2014). Here we investigate the role of another class of candidate cell swelling sensors, mechanosensitive ion channels (MS) (Cazalé et al., 1998; Haswell and Verslues, 2015; Kobayashi et al., 2018).

MS ion channels are multimeric proteins embedded in the membrane that mediate ion flux across the membrane in response to lateral membrane tension, and are found in all kingdoms of life (Martinac, 2012; Ranade et al., 2015; Basu and Haswell, 2017). Rapid cell swelling is likely to lead to an immediate increase in membrane tension, leading to the opening of MS ion channels and thereby to the exit of osmolytes and/or entry of calcium signals that regulate downstream adaptive events. In *Escherichia coli*, the MS ion channels MscS and MscL serve as osmotic safety valves, opening in response to severe hypo-osmotic shock and mediating the release of osmolytes (Levina et al., 1999; Sukharev, 2002; Belyy et al., 2010). The subsequent departure of water from the cell presumably reduces swelling and preserves cell integrity. It is also possible that *Ec*MscS mediates the swelling-induced entry of calcium, leading to intracellular signaling events (Cox et al., 2013). In plants, the overexpression of members of the MCA family leads to increased Ca^2+^ uptake in response to hypoosmotic shock in Arabidopsis, tobacco, and rice; but are not strictly required for swelling-induced Ca^2+^ uptake (Nakagawa et al., 2007; Kurusu et al., 2012b; Kurusu et al., 2012a). Members of the MscS-Like (MSL) family of MS ion channels have been implicated in osmotic homeostasis in chloroplasts (Haswell and Meyerowitz, 2006; Veley et al., 2012), and pollen (Hamilton et al., 2015), but a direct function in cell swelling perception and signal transduction has not been demonstrated.

*Arabidopsis thaliana* MSL10 is an excellent candidate for a general sensor of cell-swelling in plant cells. It is a mechanosensitive ion channel directly gated by lateral membrane tension as indicated by single-channel patch clamp electrophysiology (Haswell et al., 2008; Maksaev and Haswell, 2012). MSL10 is essentially non-selective with a slight preference for anions and has a conductance of about 100 pS. An *MSL10p::GUS* expression reporter and publicly available databases indicate that it is ubiquitously expressed at relatively low levels, with the strongest signal detected in the root and shoot apices and in the vasculature (Winter et al., 2007; Haswell et al., 2008; Laubinger et al., 2008; Zou et al., 2015).

Overexpression of *MSL10-GFP* leads to dwarfing, the induction of cell death, the hyperaccumulation of ROS, and upregulation of a suite of genes associated with cell death and ROS (Veley et al., 2014). These same phenotypes are also observed in EMS-induced gain-of-function mutant *msl10-3G*, also called *rea1* (Zou et al., 2015; Basu et al., 2020). The ability of MSL10 to induce all of these phenotypes is inhibited by the presence of phosphor-mimic lesions at seven sites in the soluble MSL10 N-terminus and is promoted by the presence of phosphor-dead lesion at these same sites (Veley et al., 2014; Basu et al., 2020). Unexpectedly, the ability to induce cell death in a heterologous system is retained in MSL10 deletion mutants lacking the predicted pore domain (Veley et al., 2014) or MSL10 with point mutants that block channel conductance (Maksaev et al., 2018). These results suggest that MSL10 has both conducting and non-conducting functions, and we have proposed that increased lateral membrane tension induces conformational changes in MSL10 that both lead to ion flux and to dephosphorylation and activation of the N-terminus (Basu et al., 2020). Our understanding of MSL10 function has been limited to these effects of MSL10 gain-of-function alleles, and its normal physiological function remains unclear. Here we describe our efforts to test the hypothesis that MSL10 serves as a sensor of membrane tension during cell swelling.

## RESULTS

We developed a seedling-based assay to determine the role played by MS ion channels in cell swelling responses. As gain-of-function *MSL10* alleles trigger cell death (Veley et al., 2014; Basu et al., 2020), we first assessed cell death after cell swelling, using Evans blue uptake. A diagram of the treatment protocol is shown in **Figure 1A**. Seedlings were germinated and grown vertically on solid media supplemented with either 140 mM mannitol or −0.5 MPa PEG (polyethylene glycol). These concentrations were selected because they allow approximately 75% germination and do not affect germination or root growth in a genotype-specific manner (**Figure S1A-D**). After 5-6 days, seedlings were transferred to liquid media containing 140 mM mannitol or −0.5 MPa PEG and either 600 nM isoxaben (ISX) or 500 nM 2,6-dichlorobenzonitrile (DCB). Both ISX and DCB are inhibitors of cellulose biosynthesis (Tateno et al., 2015). Seedlings were incubated with gentle shaking for 4 hours, then moved to solutions with or without mannitol or PEG for iso- or hypo-osmotic treatments, respectively. Hypo-osmotic treatment led to root cells that were significantly wider compared to iso-osmotic treatment, but did not lead to substantial changes in cell length nor cell bursting rates Figure S1E-H.

**Figure 1.**
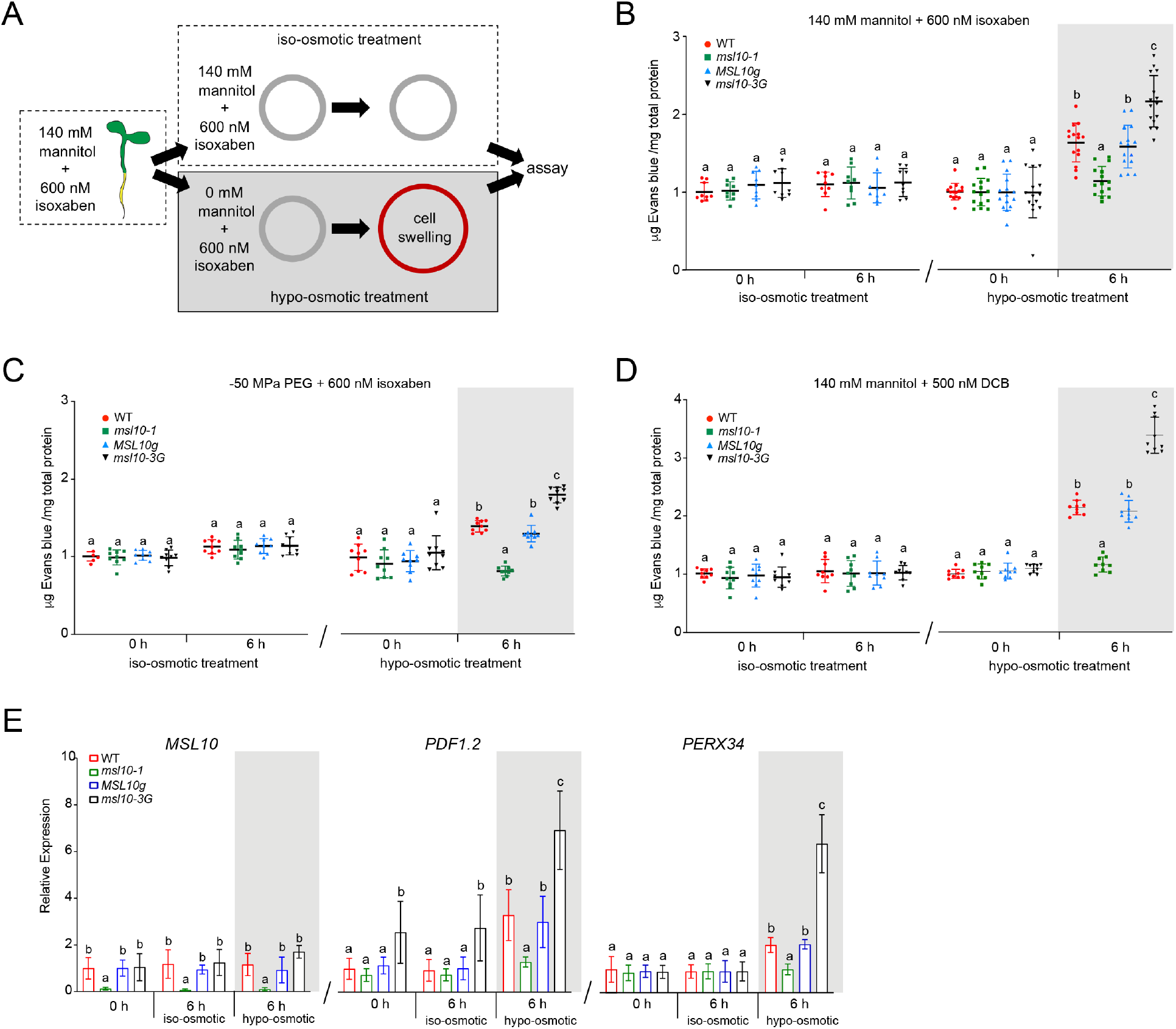
Cell swelling results in MSL10-dependent cell death. **(A)** Diagrammatic representation of the seedling cell swelling assay and the expected effects of each treatment on a single cell (indicated with a grey or red circle). **(B), (C), (D)** Quantification of cell death as assessed by Evans blue uptake, normalized to iso-osmotic treatment of the wild type. Five-day-old seedlings were incubated with either 600 nM ISX **(B and C)** or 500 nM DCB (**D)** in either 1X MS supplemented with 140 mM mannitol **(B, D)** or 1/2X MS supplemented with −0.5 MPa PEG **(C)**. After four hours incubation, seedlings were treated to either iso-osmotic or hypo-osmotic treatments. Reported values are normalized to wild type iso-osmotic treatment at 0 h. Results from five **(B)** or three **(C-D)** independent trials are shown. Each trial included three replicates of a pool of six seedlings for each genotype and treatment. **(E)** Quantitative RT-PCR analysis of *MSL10* and two genes previously shown to be upregulated in *MSL10* gain-of function lines. cDNA was synthesized from total RNA extracted from 18 five-day-old seedlings grown on 1X MS supplemented with 140 mM mannitol and subjected to cell swelling for the indicated time points. Results from three independent trials are shown. Each trial included three technical replicates of a pool of 18 seedlings for each genotype and treatment. Expression levels were normalized to both *EF1α* and *UBQ5*. Mean fold-change values relative to the wild type are plotted the mean of three biological replicates. In all experiments, error bars represent standard deviation (SD) between the means and two-way ANOVA with posthoc Tukey’s test was used to assess statistical differences; different letters denote significant differences (P < 0.05).

### Cell swelling induces cell death

In wild-type seedlings, the amount of cell death as measured by Evans blue uptake increased an average of 1.6-fold after 6 hours of hypo-osmotic treatment, but no significant increase was observed with iso-osmotic treatment (**Figure 1B and Figure S2A**). Similar results were obtained when −0.5 MPa PEG-infused agar was used in place of 140 mM mannitol (**Figure 1C**), or when DCB (a 2.2-fold increase in Evans blue uptake) was used instead of ISX (a 1.5-fold increase in Evans blue uptake). Seedlings treated with 140 mM mannitol, −0.5 MPa PEG, ISX, or DCB alone did not show an increase in cell death with hypo-osmotic treatment, though the relatively unhealthy seedlings grown on 330 mM mannitol alone did (**Figure S2B-C**). Shaking during the assay did not affect results (**Figure S2D**). Although the seedlings remain submerged for the entire duration of cell swelling assay, no increase in cell death was detected after iso-osmotic treatment, indicating that anoxia does not contribute to the observed increase in cell death seen with hypo-osmotic treatment. Collectively, these results indicate that a combination of cell wall biosynthesis inhibition and cell swelling leads to increased cell death in Arabidopsis seedlings.

### MSL10 promotes cell swelling-induced cell death

We next determined the role of MSL10 in the cell swelling-induced cell death in seedlings. As shown in **Figure 1 B-D**, MSL10 is required for cell death in response to cell swelling in all three treatment regimens (mannitol + ISX, PEG + ISX, and mannitol + DCB). First, *msl10-1* null mutants (**Figure 1B-D**) did not show any increase in cell death after 6 hours of hypo-osmotic swelling, while *msl10-1* null mutants complemented with the wild-type genomic copy of *MSL10* (*MSL10g*) had cell death levels indistinguishable from the wild-type in mannitol + ISX, PEG + ISX, and mannitol + DCB treatment regimes, respectively. Second, a heightened cell death response was observed in *msl10-3G* gain-of-function mutant lines in all three regimes (1.4-fold, 1.5-fold and 1.8-fold increases over wild type at 6 h for mannitol + ISX, PEG + ISX and mannitol + DCB, respectively). Quantitative gene expression analysis revealed no induction of *MSL10* transcripts in response to either iso-osmotic or hypo-osmotic conditions in any of the genotypes tested (**Figure 1E)**. As expected, *MSL10* transcripts were undetectable in *msl10-1* lines. The amount of cell swelling and bursting in *ms10-1* and *msl10-3G* lines were not distinguishable from the wild type (**Figure S1E-G**).

We also investigated if genes previously known to be upregulated in *MSL10* gain-of-function lines (Veley et al., 2014; Zou et al., 2015; Basu et al., 2020) were also upregulated in response to swelling-induced cell death. Wild type, *msl10-1*, and *msl10-3G* mutant plants were subjected to cell swelling and levels of *PLANT DEFENSIN1.2* (*PDF1.2*) and *PEROXIDASE34* (*PERX34*) assessed by quantitative RT-PCR. In wild type lines, *PDF1.2* levels increased 3.8-fold and *PERX34* increased 2.2-fold in response to cell swelling (**Figure 1E**). The *msl10-3G* mutant displayed an even higher expression of *PDF1.2* (5.4-fold) and *PERX34* (5-fold) in response to cell swelling, whereas no noticeable change in either of these two genes was observed in the *msl10-1* seedlings. These results show that an MSL10-dependent mechanism, likely related to the phenotypes observed in *MSL10* gain-of-function lines, leads to increased cell death in response to cell swelling.

### MSL10 potentiates a transient increase in cytoplasmic calcium in response to cell swelling

As described above, one of the most well-documented responses to hypo-osmotic shock is a transient increase in cytoplasmic calcium ([Ca^2+^]_cyt_). We therefore determined if this response was also induced in response to cell swelling and evaluated the role of MSL10. The calcium-activated luminescence of Aequorin provides a quantitative read-out of in *vivo* changes in [Ca^2+^]_cyt_ in Arabidopsis seedlings (Knight et al., 1991; Zhu et al., 2013). Wild type, *msl10-1*, and *msl10-3G* seedlings expressing Aequorin were challenged with a hypo-osmotic treatment (by diluting 1:1 in deionized water) or an iso-osmotic treatment (by diluting 1:1 in 140 mM mannitol) using the injection function of a plate reader. In wild-type seedlings, [Ca^2+^]_cyt_ rose sharply within 2 seconds of hypo-osmotic treatment (black arrow), then almost immediately declined to a level near the baseline (**Figure 2A**). In *msl10-1* null mutants this peak was greatly diminished, showing approximately the same peak height as upon iso-osmotic treatment (**Figure 2B**). Conversely, *msl10-3G* seedlings showed a much higher peak of [Ca^2+^]_cyt_ in response to cell swelling than wild-type seedlings. **Figures 2C** and **D** show comparisons of calcium levels at four points during experiments: before treatment (2.2 seconds after the start of the experiment), at the apex of the main peak (5.6 seconds after the start of the experiment), and at two points during recovery (7.8 and 10 seconds after the start of the experiment). The [Ca^2+^]_cyt_ response to hypo-osmotic treatment in the *msl10-1* was significantly lower than that in the wild type at both 5.6 and at 7.8 seconds, resembling the response to iso-osmotic treatment.

**Figure 2.**
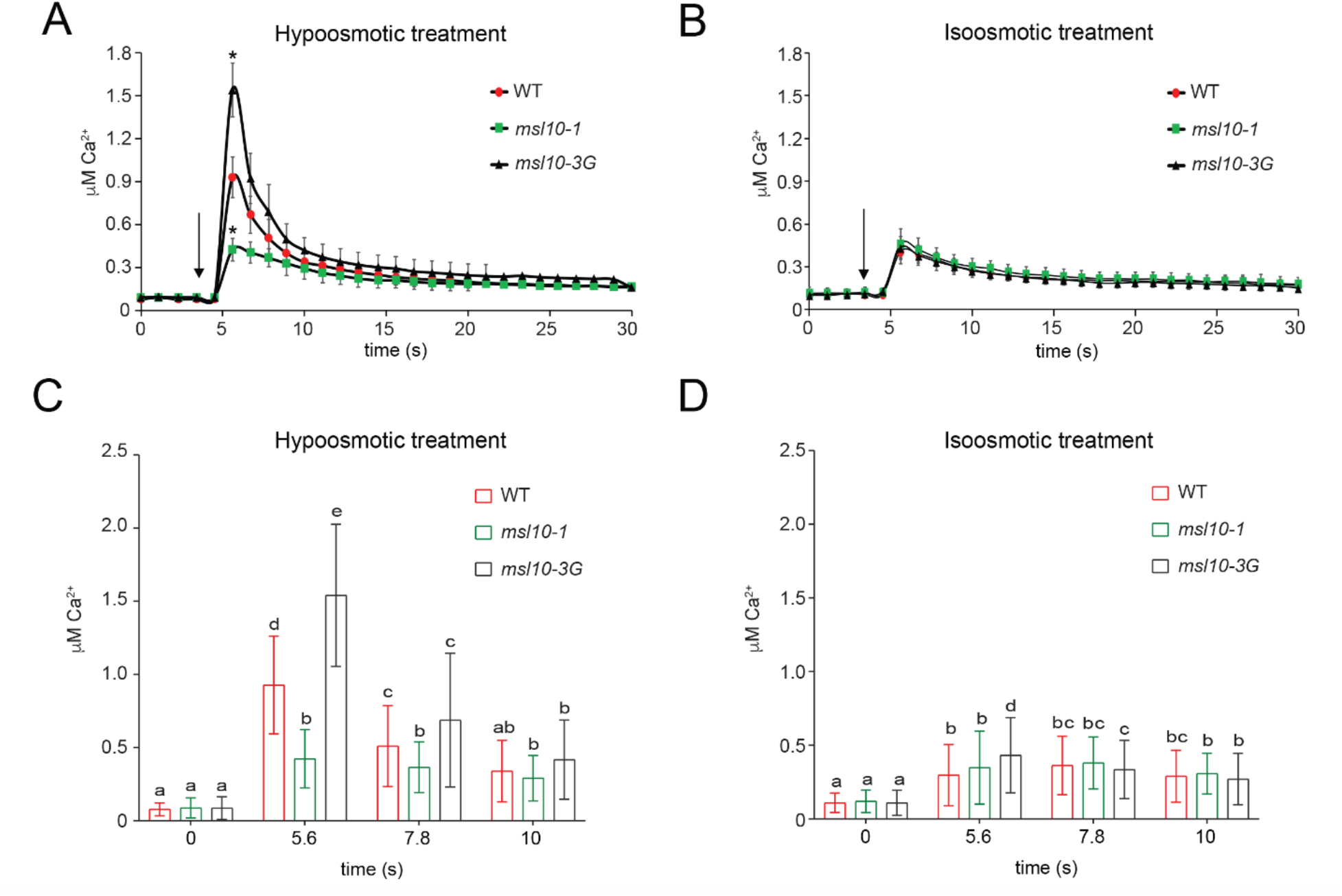
MSL10 potentiates a transient increase in cytoplasmic calcium in response to cell swelling. **(A)** and **(B)** Time courses of [Ca^2+^]_cyt_ in 3- to 4-day-old aequorin-expressing seedlings in response to hypo-osmotic (**A, C**) or iso-osmotic (**B, D**) treatment. Errors bars indicate standard error (SE) of the means from four independent trials, each including three transgenic lines per genotype, n = 7 seedlings per line. Arrows indicate application of deionized water or 140 mM mannitol to produce hypo- or iso-osmotic treatments, respectively. **(C)** and **(D)** Comparison of [Ca^2+^]_cyt_ levels during pretreatment from the data shown in (A) and (B), respectively, at 5.6, 7.8 and 10 seconds after stimulation. Error bars and statistical analyses are as in Figure 1.

The [Ca^2+^]_cyt_ response in the *msl10-3G* mutant was significantly higher than the wild type at 5.6 seconds. **Figure 2** shows compiled results of four independent trials, each of which included seedlings from three transgenic lines of each background; variation among individual transgenic lines is shown in **Figure S3 A-F**. Thus, cell swelling triggers a rapid and transient increase in [Ca^2+^]_cyt_ and MSL10 is required for this response. We note that most previous reports of [Ca^2+^]_cyt_ response to hypo-osmotic shock describe a biphasic response (Takahashi et al., 1997b; Pauly et al., 2001; Hayashi et al., 2006; Shih et al., 2014). However, these responses vary widely in timescale and relative peak heights, and in some cases a monophasic response is observed (Takahashi et al., 1997a; Kurusu et al., 2012a; Nguyen et al., 2018). These differences can be attributed to differences in the tissue used, the extent of osmotic down shock, or the timescale of measurement.

### MSL10 potentiates ROS accumulation in response to cell swelling

As described above, plant cells hyperaccumulate ROS in response to hypo-osmotic shock, ROS accumulation is commonly associated with the induction of various types of PCD in plants (Petrov et al., 2015; Locato et al., 2016). In addition, we previously documented increased ROS accumulation in plants harboring *MSL10* gain-of-function alleles (Veley et al., 2014; Basu et al., 2020). We therefore wished to determine if cell swelling triggers ROS hyper-accumulation, and to investigate the possible role of MSL10 in that process. Five-day-old seedlings were stained with the ROS indicator dye 2, 7-dichlorofluorescein diacetate (H_2_DCFDA) at the indicated time points after cell swelling and fluorescence intensities were measured using a laser scanning confocal microscope. As shown in **Figure 3A**, by 30 minutes cell swelling produced an almost 2-fold increase in H_2_DCFDA fluorescence intensity in wild-type seedlings. At this same time point, no increase in ROS levels was observed in *msl10-1* null mutants, but ROS accumulation was recovered in the *MSL10g* lines, and *msl10-3G* lines accumulated high levels of ROS (almost 4-fold increase in H_2_DCFDA fluorescence over the iso-osmotic treatment). By 60 minutes, however, ROS levels in the *msl10-1* line were no longer significantly different from those in the wild type. Iso-osmotic treatment did not induce ROS accumulation, beyond a small increase in *msl10-3G* lines.

**Figure 3.**
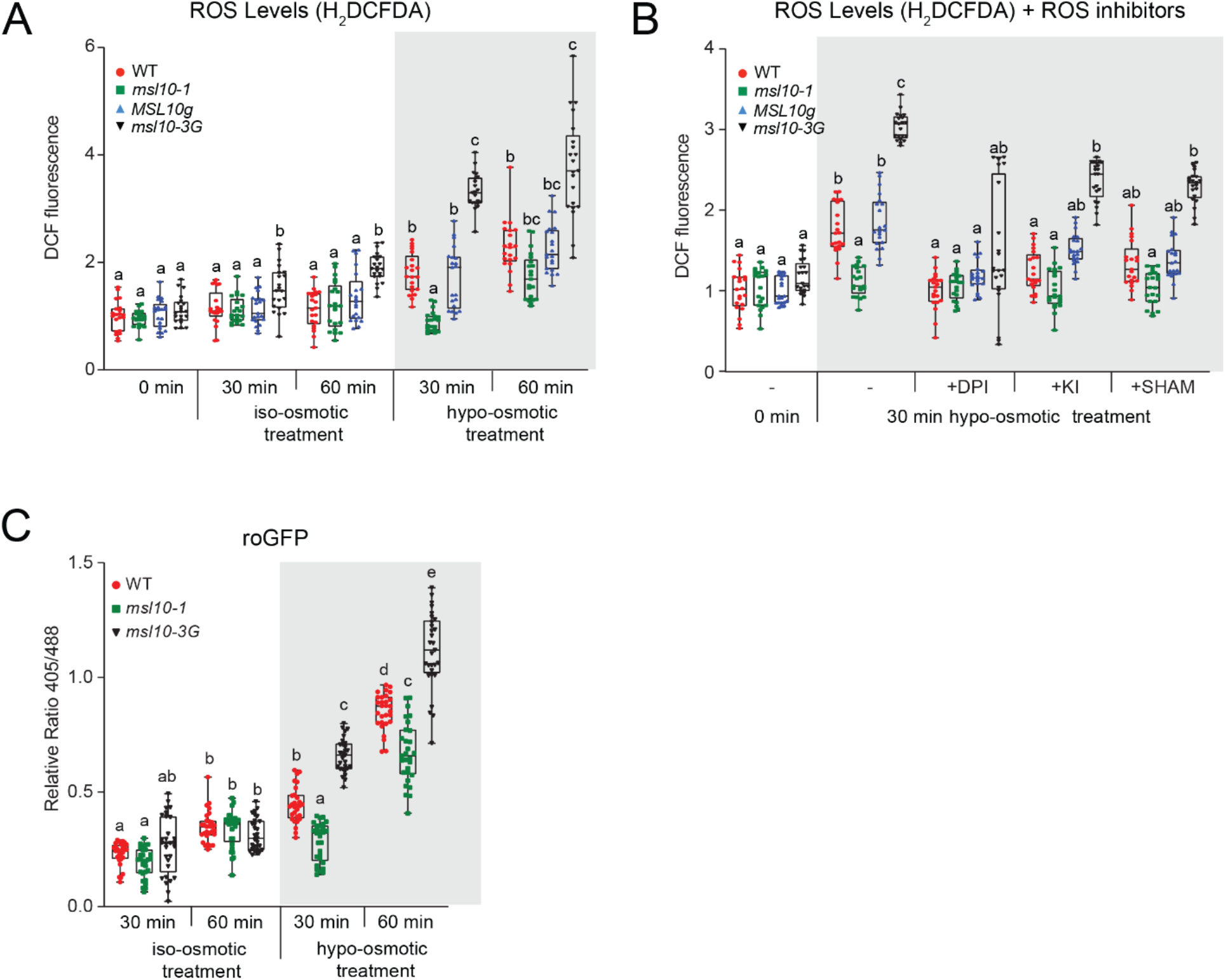
MSL10 potentiates ROS accumulation in response to cell swelling. **(A)** Quantification of H_2_DCFDA fluorescence in five-day-old seedlings in response to cell swelling. Results from three independent trials are shown. Each trial included 5-10 seedlings from each genotype and treatment. **(B)** Effect of ROS scavengers and inhibitors on ROS accumulation during cell swelling as detected by H_2_DCFDA staining. Results from three independent trials are shown. Each trial included seven seedlings for each genotype and treatment. **(C)** Relative change in roGFP1 oxidation upon cell swelling, calculated from the measured intensity ratios (405/488 nm). Five-day-old seedlings were subjected to cell swelling and fluorescent intensity ratios were quantified from confocal images of root transition/elongation zones. Results from three independent trials are shown. Each trial included 9-10 seedlings for each genotype and treatment. Error bars and statistical analyses are as in Figure 1.

We next attempted to localize the source of cell swelling-induced ROS by including inhibitors in the cell swelling treatment solutions. We tested diphenyleneiodonium (an NADPH oxidase inhibitor, DPI), potassium iodide (KI, a scavenger of H_2_O_2_) or salicylhydroxamic acid (SHAM, a peroxidase inhibitor) (Rouet et al., 2006; Dunand et al., 2007; Kurusu et al., 2012a). DPI had the strongest effect, reducing levels in the wild type and *MSL10g* lines to that of untreated controls, and also reducing levels in *msl10-3G* lines (**Figure 3B**). Including KI and SHAM in cell swelling treatments had much more modest effects on H_2_DCFDA fluorescence intensity. Thus most, but not all, of the ROS produced in response to cell swelling is dependent on MSL10, and likely involves the action of NADPH oxidases.

To further confirm the role of MSL10 in early ROS accumulation after cell swelling, we used the ROS-dependent fluorescence of cytoplasmic roGFP1, a ratiometric fluorescent biosensor that is used to report intracellular levels of ROS (Jiang et al., 2006). The redox status of roGFP1 is reflected in the its emission intensity when excited at 405 nm relative to its emission intensity when excited at 488 nm. The accumulation of ROS results in the oxidation of cysteine residues of roGFP1, and a higher 405/488 emission ratio. Wild-type, *msl10-1* and *msl10-3G* mutant lines expressing cytoplasmic roGFP1 were subjected to cell swelling followed by confocal imaging. For calibration, seedlings were treated with either 10 mM H_2_O_2_ or 10 mM DTT to fully oxidize or reduce roGFP1 (**Figure S4)**. In agreement with the H_2_DCFDA fluorescence intensity results shown in **Figure 3A**, roGFP1 signal increased in wild-type seedlings after 30 minutes of hypo-osmotic treatment (**Figure 3C**). At 30 minutes of hypo-osmotic treatment, *msl10-1* seedlings did not show any increase in roGFP1 signal, though an increase was observed by 60 minutes. In *msl10-3G* mutants, roGFP1 signal was significantly higher than the wild type at both 30 and 60 minutes after hypo-osmotic treatment. Thus, cell swelling leads to increased ROS levels within thirty minutes after treatment, and MSL10 is required for the early stage of this response. MSL10 is not strictly required for the subsequent increases in ROS levels at 60 minutes, though it appears to contribute.

### MSL10 potentiates hypo-osmotic shock-associated changes in gene expression

It was previously observed that several genes implicated in touch- and mechano-response are induced in Arabidopsis root tips after 20 minutes of hypo-osmotic treatment (Shih et al., 2014). We therefore investigated if these genes were also upregulated in our seedling assay, and if MSL10 is required for any effects. We selected 5 genes to assess by quantitative RT-PCR: *TCH1, TCH2, TCH3, TCH4*, and *WRKY18*. TCH1-3 encode calmodulin or calmodulin-like proteins. In wild-type seedlings, *TCH1, TCH2, TCH3*, and *TCH4* were upregulated 6- to 22-fold after 30 minutes of hypo-osmotic swelling, but levels returned to almost baseline (*TCH2, TCH4*), or lower than baseline (*TCH1*, *TCH3*) after 60 minutes (**Figure 4**). *WRKY18* was also upregulated, but with different temporal dynamics, showing a 4-fold increase in transcript levels after 30 minutes and a 12-fold increase after 60 minutes of hypo-osmotic treatment. MSL10 is required for full induction of all of these genes in response to hypo-osmotic swelling, as *msl10-1* null mutants exhibited about half the transcript increases compared to the wild type. In all five cases, the complemented *msl10-1* mutant, *MSL10g*, was indistinguishable from the wild type, and the gain-of-function line *msl10-G* showed an even higher increase in transcript levels than the wild type. Taken together, the results presented in **Figures 2, 3**, and **4** show that cell swelling activates a number of downstream responses previously associated with hypo-osmotic shock, including cytoplasmic calcium influx, a ROS burst, and the induction of mechano-inducible genes.

**Figure 4.**
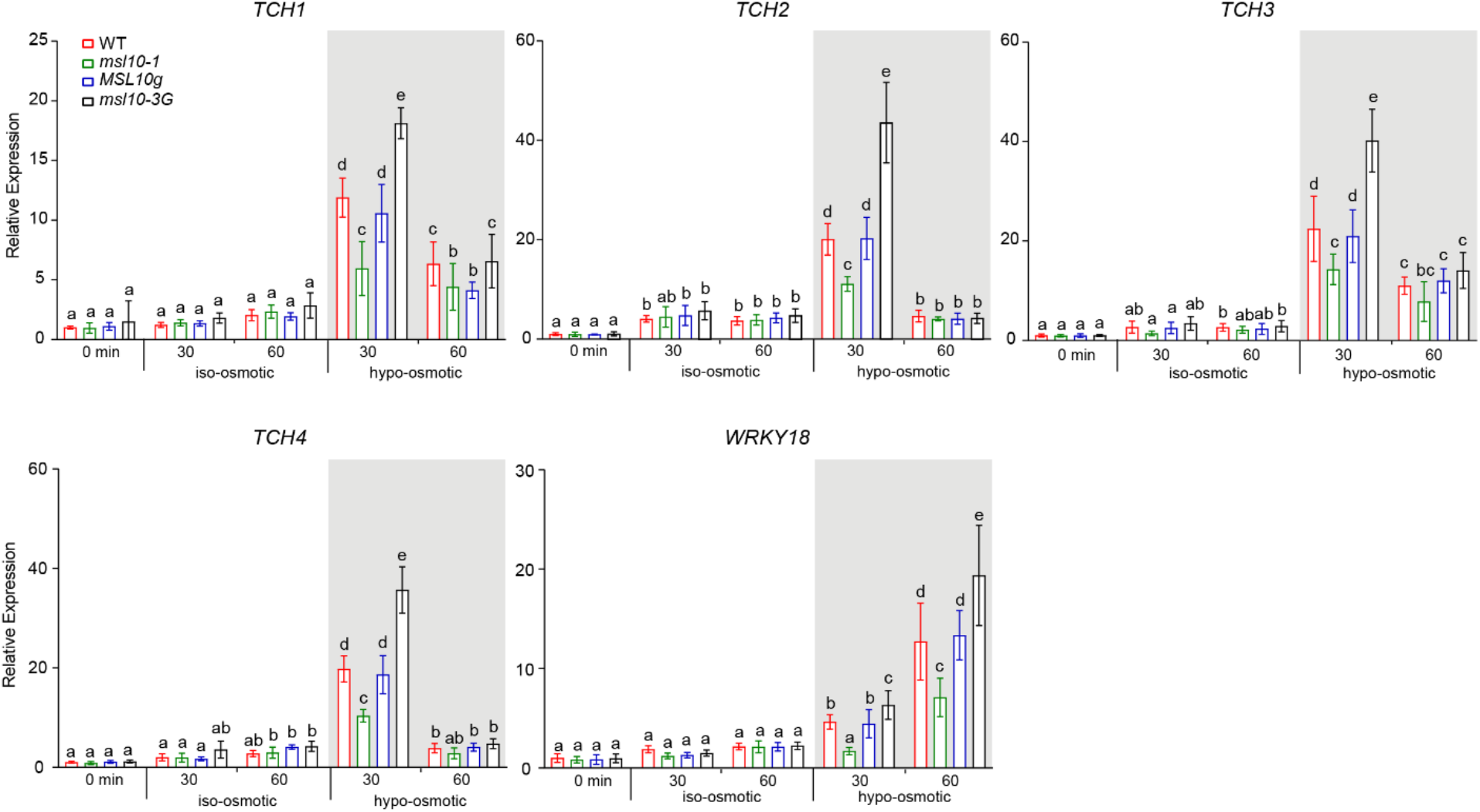
MSL10 potentiates hypo-osmotic treatment-associated gene expression. Quantitative RT-PCR analysis of genes implicated in hypo-osmotic stress response. Performed and analyzed as in Figure 1E.

### Cell swelling results in MSL10-dependent programmed cell death

Programmed cell death (PCD) is a generic term used for multiple processes that result in induced and highly regulated cellular suicide in both plants and animals (van Doorn, 2011). In plants, PCD can be generally categorized as developmentally-induced (dPCD), pathogen-induced (pPCD) or environmentally-induced (ePCD) (Olvera-Carrillo et al., 2015; Kumar et al., 2016; Huysmans et al., 2017). As identifying and categorizing PCD-like events in plants has proven difficult and controversial (van Doorn, 2011; de Pinto et al., 2012), we used a set of well-established assays for PCD, including the terminal deoxynucleotidyl transferase dUTP nick end labeling (TUNEL) assay (Tripathi et al., 2017); the activation of cysteine proteinases (caspase-like proteases) (Xu and Zhang, 2009); the upregulation of marker genes (Olvera-Carrillo et al., 2015); and acidification of the cytosol (Young et al., 2010) .

First, we used the TUNEL assay for detecting DNA fragmentation in root cells of seedlings subjected to iso-osmotic or hypo-osmotic treatments (**Figure 5A and Figure S5A**). After 6 hours of cell swelling, the number of TUNEL-positive nuclei doubled in wild-type seedlings. Furthermore, this increase required MSL10, as no increase over iso-osmotic treatment was observed in *msl10-1* mutants, and levels were restored in the *msl10-1; MSL10g* line. The *msl10-3G* gain-of-function line showed even higher levels of cell swelling-induced TUNEL staining than the wild type (a 5-fold increase after 6 hours of cell swelling). A small increase in DNA fragmentation in the *msl10-3G* line above basal levels was observed. As expected, essentially all cells had TUNEL-positive nuclei when wild type seedlings were incubated with DNase I for 15 min prior to the TUNEL reaction, and no TUNEL-positive nuclei were detected when the terminal deoxynucleotidyl transferase enzyme was omitted from the TUNEL reaction mixture (**Figure S5B-C**).

**Figure 5.**
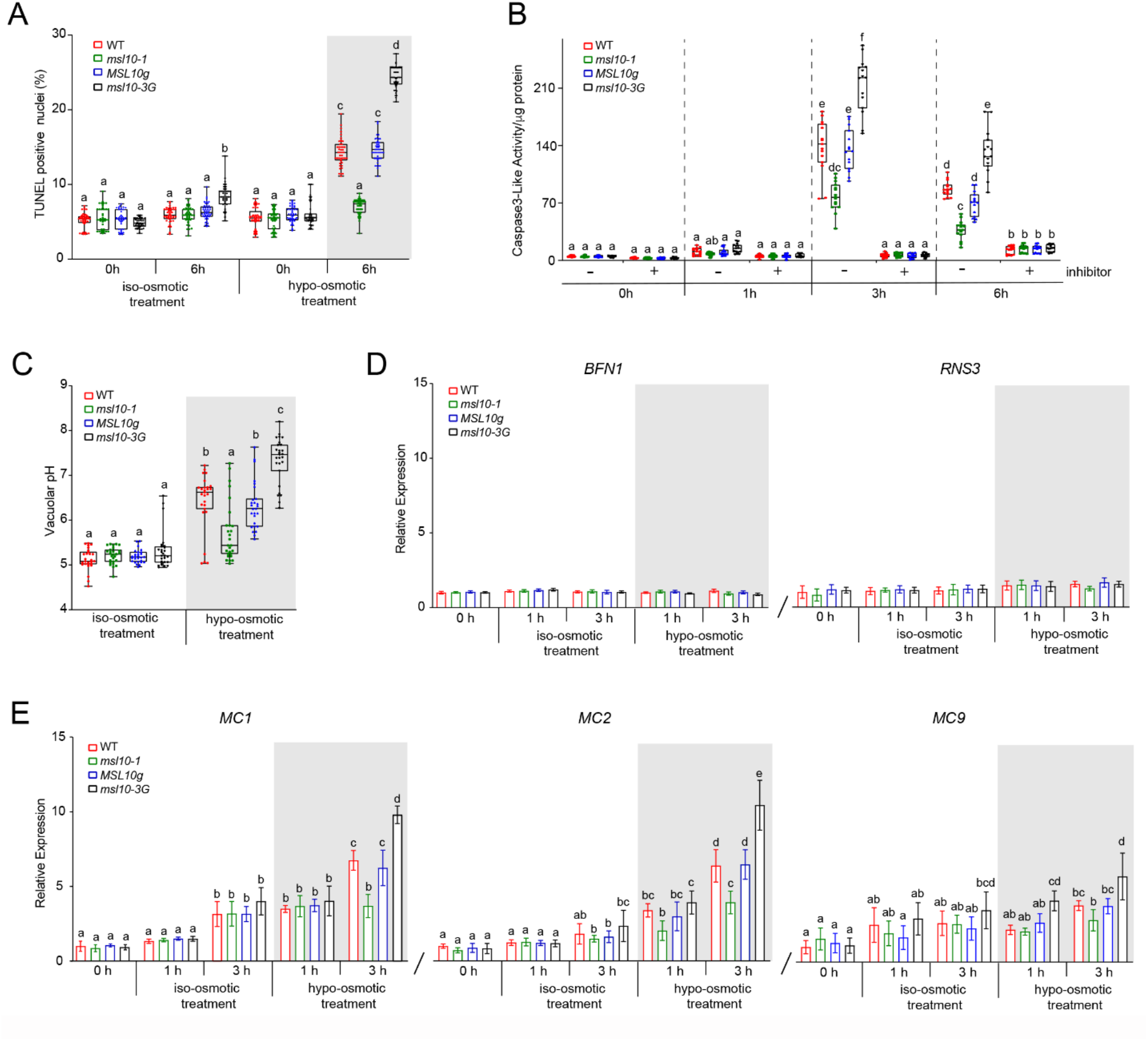
Cell swelling results in MSL10-dependent programmed cell death. **(A)** Frequency of TUNEL-positive nuclei in the root tip of seedlings after iso-osmotic or hypo-osmotic treatment. Results from three independent trials of 15 seedlings from each genotype and treatment are presented. **(B)** Caspase 3-like activity in response to cell swelling. Homogenates isolated from seven-day-old seedlings that underwent cell swelling were incubated with 50 μM Ac-DEVD-AMC for 1h. Some extracts were treated with 100 μM Ac-DEVD-CHO for 1 h before the assay. Results from three independent trials are shown. Each trial included five replicates of a pool of six seedlings for each genotype and treatment. **(C)** Increase in vacuolar pH in response to cell swelling as indicated by the pH-sensitive fluorescent dye BCECF-AM (10 μM). Results from three independent trials of 9 seedlings from each genotype and treatment are presented. **(D, E)** Quantitative RT-PCR analysis of selected genes, performed as for Figure 1E. For all panels, error bars and statistical analyses are as in Figure 1.

Next, we used a commercial fluorometric caspase-3 activity assay to assess the induction of protease activity in protein extracts from whole seedlings. This approach has been previously used in plant cells (Danon et al., 2004; Ge et al., 2016). Wild-type and complemented *MSL10g* seedlings displayed increased caspase-3-like protease activity after cell swelling (from 5 to 140 and 134 units/μg of total protein after 3 hours respectively) (**Figure 5B**). We observed a significantly smaller induction of caspase-3-like activity in *msl10-1* mutant seedlings and a significantly larger increase in the *msl10-3G* mutant in response to cell swelling, compared to the wild type. At 6 hours, a decrease in swelling-induced caspase-3-like activity was observed in all genotypes. Extracts from 1 h, 3 h, and 6 h samples that were treated with the caspase-3 inhibitor Ac-DEVD-CHO were indistinguishable from the 0-hour timepoint, indicating that the activity detected is specific to the caspase-3-like protease family.

Cytoplasmic acidification is another cellular marker of plant PCD. An increase in the typically low vacuolar pH can be used as a proxy for cytoplasmic acidification (Roberts et al., 1984; Wilkins et al., 2015). We stained seedlings with the ratiometric pH indicator dye 2,7-bis-(2-carboxyethyl)-5-(and-6)-carboxy-fluorescein acetoxy-methyl ester (BCECF-AM), which accumulates in the vacuole (Bassil et al., 2013)(**Figure 5C**). After 3 hours of cell swelling, BCECF-AM-stained root cells were visualized using confocal microscopy. The ratio of emission after excitation at 488 nm to emission after excitation at 445 nm is positively correlated with pH. The ratio of emission intensities from the two BCECF-AM excitation wavelengths was translated to vacuolar pH using an *in situ* standard curve (**Figure S5D)**. None of the genotypes displayed any statistically significant changes in vacuolar pH during iso-osmotic treatment. By contrast, cell swelling led to an increase in the vacuolar pH of wild type seedling root cells. Consistent with the TUNEL and capase-3-like activity assays, *msl10-3G* mutant seedlings displayed a greater increase in vacuolar pH in response to cell swelling than wild-type seedlings, whereas no significant increase in vacuolar pH was observed in *msl10-1* mutant seedlings.

We next turned to marker gene expression to understand what type of PCD is induced by cell swelling. *BIFUNCTIONAL NUCLEASE I (BFN1), RIBONUCLEASE 3 (RNS3)*, and *METACASPASE 9 (MC9)* are three of the nine genes belonging to the core “developmental cluster” known to be induced during developmental PCD, but not during environmentally triggered PCD (Olvera-Carrillo et al., 2015); though we note that an earlier report on *MC9* indicates that it is also involved in biotic stress (Watanabe and Lam, 2011). Since cell swelling induced detectable DNA fragmentation by 6 hours (**Figure 5A**), we decided to assess gene expression at earlier time points. *BFN1* and *RNS3* expression levels did not change significantly in response to iso-osmotic treatment or to cell swelling in any of the genotypes at any time points tested (**Figure 5D**). Only a slight increase in *MC9* levels was observed in *msl10-3G* seedlings after at 3 hours of cell swelling (**Figure 5E**). These data indicate that cell swelling-induced PCD is distinct from known developmental PCD.

Finally, we measured transcript levels of *METACASPASE 1 (MC1)* and *METACASPASE 2 (MC2)* in response to cell swelling. *MC1* is a canonical pathogenesis-related PCD metacaspase gene and *MC2* has been implicated in PCD associated with biotic stresses (Watanabe and Lam, 2011; Coll et al., 2014; Yao et al., 2020). In wild type seedlings, *MC1* and *MC2* transcript levels increased 4-6-fold after 1 hour and 3 hours of cell swelling, respectively (**Figure 5E**). In contrast, *msl10-1* null mutants only displayed a slight increase in expression of *MC2*, whereas *msl10-3G* displayed a marked increase in *MC1* and *MC2* expression (10-fold and 10.5-fold, respectively).*MC1* and *MC2* transcript levels did increase after 3 hours of iso-osmotic treatment in all genotypes, but not to the same extent as with cell swelling. Taken together, the data presented in **Figure 5** show that cell swelling induces PCD that is distinct from developmentally induced PCD, and that these effects either require or are promoted by the mechanosensitive ion channel MSL10.

### Cell swelling-induced PCD is prevented by phosphomimetic amino acid substitutions in the MSL10 N-terminus

We previously showed that overexpressed MSL10-GFP loses the ability to trigger cell death when four residues in its soluble N-terminal domain are replaced with phosphomimetic residues (MSL10 S57D, S128D, S131E, and T136D, denoted as MSL10^4D^-GFP) (Veley et al., 2014). Conversely, replacing these four residues along with three others in the N-terminus with alanine (MSL10^7A^-GFP) produces constitutive cell death in a transient expression system (Veley et al., 2014). We therefore decided to examine the effect of these same N-terminal substitutions on cell swelling-induced PCD.

Lines overexpressing *MSL10-GFP* from the strong, constitutive 35S promoter showed significantly more swelling-induced cell death than the wild type (compare 3.5-fold and 1.7-fold increase in Evans blue uptake and 21% versus 11% TUNEL-positive nuclei, respectively) (**Figure 6A, C**). However, lines constitutively overexpressing *MSL10^4D^-GFP* at high levels did not induce PCD in response to cell swelling. This cannot be attributed to differences in protein levels, as the anti-GFP immunoblot shown in **Figure 6B** indicates that MSL10-GFP and MSL10^4D^-GFP protein accumulated to similar levels in these lines. It was not possible to isolate stably transformed lines expressing phospho-dead versions of MSL10-GFP from the 35S promoter (Veley et al., 2014).

**Figure 6.**
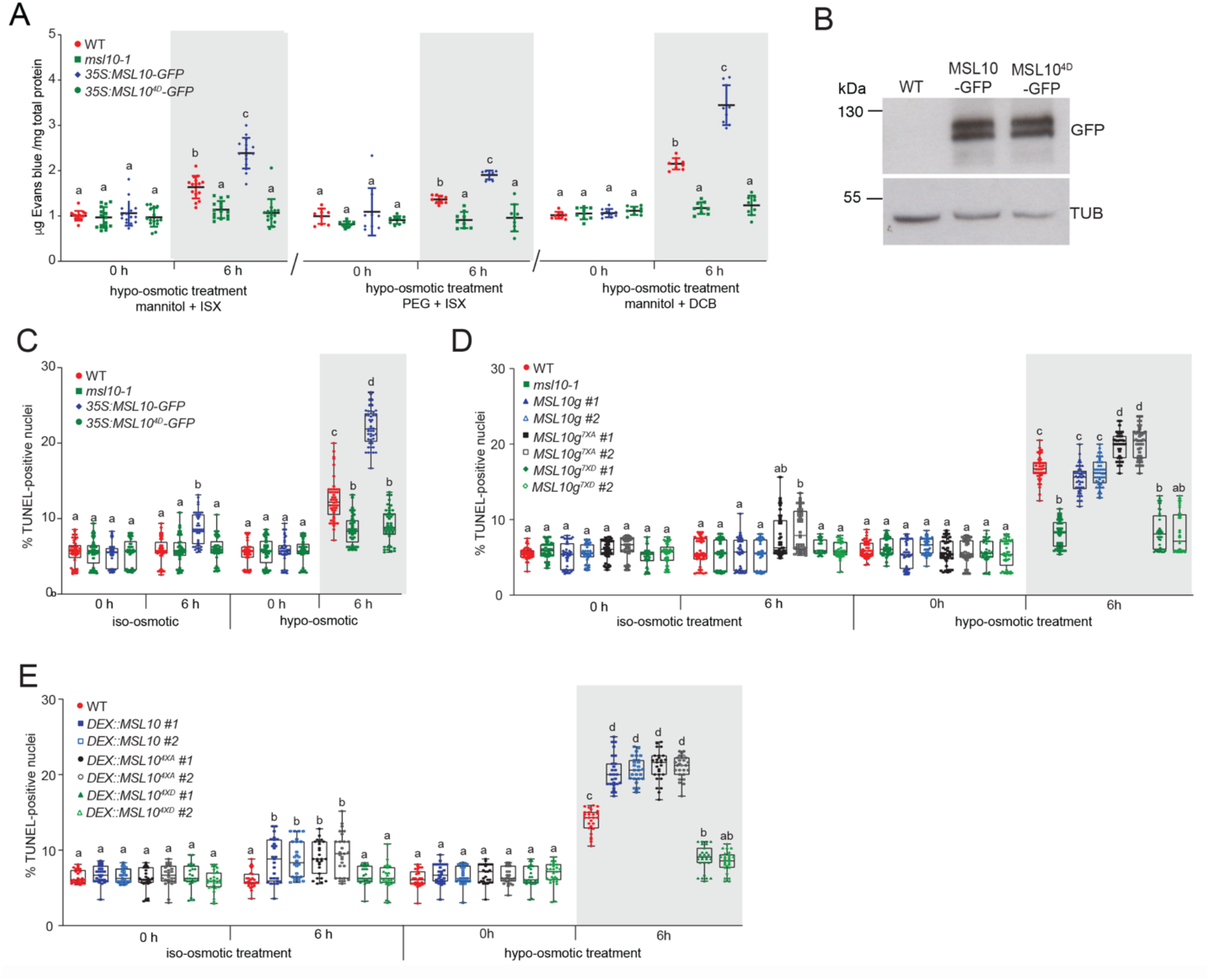
Cell swelling-induced PCD is prevented by phosphomimetic amino acid substitutions in the MSL10 N-terminus. **(A)** Swelling-induced cell death in lines over-expressing MSL10-GFP and MSL10^4D^-GFP, as measured by Evans blue uptake as described in **Figure 1**. Data are normalized against wild type at 0 h. Results from five independent trials are shown. Each trial included three replicates of a pool of six seedlings for each genotype and treatment. **(B)** Immunodetection of MSL10-GFP variant proteins. Total protein was isolated from 6 pooled five-day-old seedlings grown on 1 X MS supplemented with 140 mM mannitol. Anti-GFP primary was used to detect MSL10-GFP (top panel), and the blot re-probed with an anti-α-tubulin primary antibody (bottom panel). Theoretical molecular masses of proteins are indicated at the left according to a commercially available standard. Presence of two bands may be attributed to posttranslational modification. **(C)** *In situ* detection of DNA fragmentation in in lines over-expressing MSL10-GFP and MSL10^4D^-GFP using the Click-iT TUNEL Alexa Fluor 647 Imaging Assay kit. Results from three independent trials are shown. Each trial included 15 seedlings from each genotype and treatment. **(D)** Detection of *in situ* DNA fragmentation in response to cell swelling in seedlings expressing *MSL10g, MSL10g^7A^*, or *MSL10g^7D^* in the *msl10-1* background, as described for **Figure 5A**. Results from three independent trials are shown. Each trial included 10-20 seedlings from each genotype and treatment. **(E)** Detection of *in situ* DNA fragmentation in response to cell swelling wild type seedlings expressing *DEX:MSL10, DEX:MSL10^4A^, DEX:MSL10^4D^*, as described for **Figure 5A**. Seedlings were incubated with 30 μM DEX in 1X MS media supplemented with 140 mM mannitol during the equilibration step. Results from three independent trials are shown. Each trial included 9 seedlings from each genotype and treatment.

Furthermore, expressing untagged *MSL10* from its endogenous promoter in the *msl10-1* mutant background both produced cell swelling-induced PCD at levels comparable to the wild type (an average of 14.8% compared to 15.1%; **Figure 6D)**, and conditional overexpression of untagged *MSL10* from a dexamethasone-inducible promoter produced significantly more PCD in response to cell swelling than the wild type (an average of 20.2% compared to 14%; **Figure 6E**). However, neither expressing *MSL10^7D^* at endogenous levels nor induced expression of *MSL10^4D^* induced PCD in response to cell swelling, and these lines were statistically indistinguishable from the *msl10-1* null line. On the other hand, endogenous expression of *10gMSL10^7A^* or conditional expression of phospho-dead *MSL10^4A^* resulted in even more cell swelling-induced PCD than wild-type seedlings (20% and 21.2%, respectively, compared to an average of 15% for the wild type). We were unable to assess MSL10 protein levels in these lines as MSL10 was untagged, but transcripts from the *10gMSL10^7D^* and *10gMSL10^7D^* transgenes were similar to levels from the endogenous locus in the wild type background (**Figure S6A, S6B**).

Thus, introducing phosphomimetic lesions into the MSL10 N-terminus prevented PCD in response to cell swelling--and are consistent with the proposal that dephosphorylation of the MSL10 N-terminus is required for cell swelling-induced PCD. The observed promotion of cell swelling-induced PCD by MSL10 cannot be attributed to the presence of a GFP tag, to non-specific effects of MSL10 overexpression, nor to a developmental response to the presence of high levels of MSL10.

## DISCUSSION

Here we establish and validate a procedure for imposing cell swelling on whole seedlings. We show that cell swelling induces programmed cell death, ROS accumulation, the elevation of cytoplasmic Ca^2+^ concentration and the expression of mechano-inducible genes. Furthermore, we show that MSL10 positively influences all these steps in the swelling-induced signaling pathway, and that this effect is regulated by phosphorylation of the soluble MSL10 N-terminus.

### Cell swelling in response to cell wall softening and hypo-osmotic treatment

Plant cell mechanics are primarily determined by the dynamic interplay between the cell wall and turgor pressure. The plant cell wall is a complex arrangement of cellulose microfibrils, hemicelluloses, pectins, and glycoproteins that has incredible tensile strength, yet can be finely tuned in order to generate cell shape, allow growth, and protect cell integrity (Cosgrove, 2018). While the directionality of any changes in cell volume is controlled by the orientation of cellulose microfibrils and the local stiffness of pectin, the driving force is turgor pressure, the pressure that the protoplast exerts on the cell wall (up to multiple atmospheres) (Bou Daher et al., 2018). Importantly, the plasma membrane (and therefore, any embedded mechanosensitive ion channels) sit at the interface between cell wall strength and turgor pressure. Under normal conditions, turgor pressure and cell wall strength are in balance, but when the cell wall softens or turgor increases rapidly, cell swelling occurs and membrane tension increases.

In this study we used a combination of cell wall softening (accomplished by treatment with the cellulose synthase inhibitor ISX) and increased turgor (accomplished by transferring plants to a hypo-osmotic solution) to activate downstream cell swelling signaling pathways and thereby provide a tool for dissection of the process (**Figure 1A**). While inhibition of cell wall biosynthesis has been reported to induce cell death after 10-24 hours of treatment (Duval et al., 2005; Oiwa et al., 2013; Engelsdorf et al., 2018; Zhou et al., 2020), we did not observe any increased cell death after 6 hours of treatment with ISX or DCB alone in our assay (**Figure S2B**). Thus, at least at these early timepoints, both cell wall softening, and hypo-osmotic treatment are required for the responses reported here. Furthermore, our results can be recapitulated without cell wall softening if a larger osmotic down shock is used, presumably because it produces the same level of cell swelling (**Figure S2C**).

### MSL10 as a cell-swelling sensor

Several lines of evidence support a model wherein MSL10 is the primary plasma membrane-based sensor of cell swelling. MSL10 influences all previously documented downstream effects of hypo-osmotic shock that we tested. Cell death (**Figure 1**), a burst of [Ca^2+^]_cyt_ (**Figure 2**), hyper-accumulation of ROS (**Figure 3**), and increased transcript levels for four mechano-inducible genes (**Figure 4**) all were reduced or absent in the *msl10-1* null mutant and enhanced in the *msl10-3G* gain-of-function line. In addition, four distinct hallmarks of programmed cell death (DNA fragmentation, increase in caspase-like activity, intracellular pH, and PCD-associated gene expression) increased in response to cell death in a MSL10-dependent manner. Particularly suggestive is the observation that the transient increase in [Ca^2+^]_cyt_ levels observed in *msl10-1* plants in response to hypo-osmotic treatment is indistinguishable from the response to iso-osmotic treatment. The latter is likely to be a mechanical response to the injection of the treatment solution. These data indicate that the perceptive events that occur within the first 2 second of swelling require MSL10; the immediacy of this effect and its complete requirement for MSL10 provide strong evidence that MSL10 directly receives a swelling signal at the membrane to potentiate downstream effects.

Notably, this is the first reported loss-of-function phenotype for *MSL10* other than a modest, and presumably indirect, defect in starch degradation (Fusari et al., 2017). Despite a decade of research, plants lacking functional MSL10 have performed identically to the wild type in innumerable developmental and mechanical stress assays (Haswell et al., 2008; Shih et al., 2014; Stephan et al., 2016; Tran et al., 2017; Marhava et al., 2019; Van Moerkercke et al., 2019). Until now, genetic evidence for MSL10 function has been derived solely from the phenotypes associated with MSL10 gain-of-function alleles: the overexpression of *MSL10-GFP*, the expression of untagged *MSL10^7A^* from the endogenous promoter, or the *msl10-3G (rea1)* allele (Veley et al., 2014; Zou et al., 2015; Basu et al., 2020). In these backgrounds, phenotypic effects such as dwarfing and ectopic cell death are apparent in adult plants without added stress, though seedlings are indistinguishable from the wild type. In the experiments presented here, the *msl10-3G* and *MSL10^7A^* gain-of-function lines did not differ from the wild type in that they still required cell swelling to produce cell death, PCD, [Ca^2+^]_cyt_ transients, ROS accumulation, or gene expression; however they did eventually have a stronger response than the wild type. Thus, signaling in response to cell swelling is likely to be the normal function of the MSL10 protein, and not the result of toxic or non-physiological effects.

We previously used single channel patch clamp electrophysiology to demonstrate that MSL10 channel opens in response to increased lateral membrane tension, and that this does not require association with other cellular components (Maksaev and Haswell, 2012). Thus, one straightforward interpretation of the data presented here is that cell swelling increases membrane tension, which opens the MSL10 channel pore and provides an ionic signal that activates the rest of the downstream pathway. However, MSL10 has a preference for anions, and electrophysiological experiments with large ions suggest that it does not conduct Ca^2+^ at all (Haswell et al., 2008; Maksaev and Haswell, 2012; Guerringue et al., 2018). Instead, MSL10 is more likely to releases Cl^-^ from the cell upon opening. As MSL10 has a relatively long closing time (Maksaev and Haswell, 2012), one possibility is that an extended release of Cl^-^ leads to depolarization of the plasma membrane and the activation of depolarization-activated Ca^2+^ channels (Hedrich, 2012). One alternative explanation is that tension-induced conformational changes in MSL10 lead to dephosphorylation of the N-terminus, which in turn allows it to participate in a protein-protein interaction that facilitates the activity of a Ca^2+^ channel, potentiating increased [Ca^2+^]_cyt_ and the other downstream events and eventually leading to PCD.

Here we show that seedling cell swelling induces a type of PCD that is distinct from developmentally regulated PCD and similar to that induced by other abiotic stresses such as heat, salt stress, flooding, or dehydration (Kumar et al., 2016). The induction and execution of PCD in plants, as in animals, involves a huge number of molecular signals and communication between multiple subcellular compartments and organelles (van Doorn, 2011). As a result, we employed four distinct assays to demonstrate that swelling-induced cell death is a type of PCD (**Figure 1** and **5**). These different assays produced qualitatively different results with respect to a requirement for MSL10. Cell swelling-induced increases in TUNEL-positive nuclei and cytoplasmic acidification were fully MSL10-dependent, while increased caspase-3-like protease activity and the increased expression of metacaspase genes *MC1* and *MC2* were positively influenced by, but not fully dependent on, MSL10. These differences may reflect tissue-specific differences in swelling-induced PCD as the TUNEL assay and cytoplasmic acidification assays were performed on cells of the elongation zone of roots, while the caspase-3 activity and gene expression assays were performed on whole seedlings. Based on these results, we categorize MSL10-dependent cell swelling-induced PCD as a type of ePCD.

### What, if any, is the adaptive advantage of responding to cell swelling by inducing PCD?

In the case of pathogenesis, cell suicide local to the infection is thought to prevent the growth and thus the spread of a biotrophic pathogen (Mukhtar et al., 2016). Abiotic stress-induced PCD in roots is thought to promote the development of lateral roots better acclimated to salinity (Huh et al., 2002) and to produce root architectures that are more drought- or flooding-tolerant (Subbaiah and Sachs, 2003; Duan et al., 2010). The same might be true of roots cells that have undergone swelling in our assay. A more general interpretation is that when cells are likely to have experienced membrane damage due to extreme cell swelling, triggering PCD stops growth in damaged tissues, removes damaged cells, and allows the plant to recover materials (Locato and De Gara, 2018).

The results presented here provide a physiological context for MSL10 function, link disparate observations regarding hypo-osmotic shock response in plant cells, and build a foundation for molecular exploration of the signaling pathway. We anticipate that future work will reveal how MSL10 functions as a membrane-based sensor that connects the perception of cell swelling to the downstream signaling cascade of abiotic-stress induced PCD.

## METHODS

### Plant materials

All *Arabidopsis thaliana* plants used in this study were in the Columbia-0 ecotype background. *msl10-3G* is an EMS-induced point mutant originally isolated as recessive gain-of-function mutant of *MSL10* (Zou et al., 2015). T-DNA insertional mutant seeds of *msl10-1 (SALK_07625)* obtained from the Arabidopsis Stock Centre (Haswell et al., 2008). Transgenic lines used in this study were previously characterized, including *35S:MSL10-GFP* (line *12-3*), *35S:MSL10^4D^-GFP* (line *6-15*) (Veley et al., 2014), as well as *DEX:MSL10* (line *1-2* and *4-3*), *DEX MSL10 ^4A^* (line *1-4* and *4-2*) *DEX MSL10^4D^* (line *8-3* and *2-1*) and genomic versions of *MSL10g (line 1-2* and *5-1)*, *MSL10g^7A^* (line*21-1* and *13-6*), *MSL10g^7D^* (line *17-1* and *4-22*) (Basu et al., 2020). Unless stated otherwise, all chemical reagents were obtained from Sigma-Aldrich. Seeds were surface sterilized with 50% (v/v) bleach for 5 min, washed five times with sterilized distilled, deionized water. Arabidopsis plants were germinated and grown on vertically oriented petri plates containing MS salt and 0.8% agar medium for 3-7 d depending on the experiments. For conditional induction of *MSL10*, seedlings were incubated with 30 μM dexamethasone (DEX) dissolved in 0.016% ethanol for 12 h.

### Generation of transgenic plants

In all cases, transgenes were introduced into plants using the floral dip method (Clough and Bent, 1998). For genotyping, genomic DNA was extracted from leaves as previously described (Edwards et al., 1991). Primers used for cloning and genotyping are listed in **Supplemental Table 1**. Wild type, *msl10-1* and *msl10-3G* lines were transformed with *35Sp::roGFP1* (Jiang et al., 2006). Positive transformants were selected on solid MS medium supplemented with 50 mg/L hygromycin. The T3 homozygous progenies were used for further studies.

To obtain transgenic lines expressing cytoplasmically localized apoaequorin, wild type, *msl10-1* and *msl10-3G* lines were transformed with pBINU-CYA (*YFP*-fused apoaequorin (*AEQ*) driven by an ubiquitin (*UBI10*) promoter) as previously described (Mehlmer et al., 2012). Phosphinothricin-resistant primary transformants were screened for GFP fluorescence and three independent T3 transgenic lines were used for further analysis.

### Dose-dependent sensitivity of mannitol and PEG on seed germination

Seeds were sterilized and plated on MS medium (MS salts, 0.5 g/L MES, pH 5.7, and 0.8% [w/v] agar) supplemented with the indicated concentrations of mannitol (0-500 mM) and polyethylene glycol 8000 PEG (0 to −1.2 MPa) as described (van der Weele et al., 2000; Verslues et al., 2006). Plates were stratified in darkness for 3 d at 4°C and transferred to a culture room set at 23°C with continuous light intensity of 120 μmol/m^2^/s (fluorescence). Germination (radicle emergence) percentages were determined after 5 d from the end of stratification.

### Relative root elongation

The primary root length was measured after 7 d of growth in either MS media supplemented with mannitol (0 or 140 mM mannitol) or PEG (0 to −0.5 MPa). Fifteen to nineteen seedlings per line were measured. Plates were scanned using the Syngene PXi imagining system equipped with GeneSys image acquisition software, and root length was measured using the ImageJ software.

### Cell swelling treatments

This assay was modified from (Droillard et al., 2004). Briefly, seedlings were vertically grown on either MS agar (0.8% w/v) plates supplemented with 140 mM mannitol or PEG infused (−0.5 MPa) ½X MS agar (1.5% w/v) plates for 5-6 days post germination. Seedlings were then equilibrated in liquid MS media supplemented with 140 mM mannitol or ½X MS supplemented with PEG (−0.5 MPa) for 4 h in 12-welled plates with gentle shaking. Six seedlings were placed in each well containing 2 mL of media. Either ISX (600 nM) or DCB (500 nM) was added to the liquid media during equilibration. For hypo-osmotic treatments, the existing media was replaced by MS media containing 600 nM ISX but without mannitol or PEG for 6 h and 12h. For iso-osmotic treatments, the existing media was replaced by fresh MS media supplemented with 140 mM mannitol or ½X MS supplemented with PEG (−0.5 MPa) and either ISX or DCB was added.

### Evans blue quantification

Seedlings were treated with 0.25% (w/v) Evans blue for 15 min, washed thoroughly, and homogenized in 10% methanol and 0.03% SDS. Blue color was quantified in 96-well plates in a plate reader (Infinite 200 PRO [Tecan]) by measuring absorbance at 630 nm and normalized to total protein, determined using the Quick Start Bradford Protein Assay (Bio-Rad) following manufacturer’s instruction.

### TUNEL assay

The TUNEL (terminal deoxynucleotidyl transferase-mediated dUTP nick-end labeling) reaction was used to analyze cell swelling-induced DNA fragmentation as described previously (Tripathi et al., 2017) with minor modifications. Fixation, permeabilization and washing steps were performed in 12-welled plates, and the labeling reaction and staining were performed in 1.5 mL microcentrifuge tubes. Both iso- and hypo-osmotically-treated 5-day-old seedlings were fixed in 4% paraformaldehyde in PBS, pH 7.4 for 24 h at 4°C, then washed with 100% ethanol for 10 min. After fixation, seedlings were transferred to 70% ethanol for at least 16 h at −4°C. Dehydrated seedlings were washed five times with PBS and permeabilized for 30 min at 37°C with 0.1% Triton X-100 in 100 mM sodium citrate buffer pH 6. Next, seedlings were incubated in 10 mM Tris-Cl, pH 7.5 supplemented with 20 μg/mL proteinase K for 30 min at 37°C and washed five times with PBS. In situ nick-end labeling of nuclear DNA fragmentation was performed for 1 h in the dark at 37°C using the In-Situ cell death detection kit (Roche Applied Science) according to the manufacturer’s manual. For GFP-tagged overexpression lines, the Click-it TUNEL Alexa 647 (Thermofisher) kit was used, following the manufacturer’s instruction. After TUNEL reaction seedlings were rinsed 3 times in PBS and counterstained with DAPI (2.5 μg/mL) for 30 min in the dark. After washing, seedlings were mounted in the anti-fading reagent Citifluor. For each biological replicate, a negative control omitting addition of the terminal-deoxynucleotidyl-transferase and a positive control (DNase I; 1 μg/mL) treatment was included to ensure the reproducibility of the assay.

### Caspase-3-like Activity

Seedlings were harvested at the indicated time points after cell swelling. Total cytoplasmic protein is extracted by homogenizing seedling to a powder in liquid nitrogen with a mortar and pestle. The powder was collected in 1.5 ml tubes and resuspended in assay buffer (20% glycerol, 0.1% Triton, 10 mm EDTA, 3 mM dithiothreitol, 2 mm phenylmethylsulfonyl fluoride, and 50 mM sodium acetate, pH 7.4) as described (Danon et al., 2004). Caspase-3-like activity was assessed using a commercial kit (Caspase 3 Fluorometric Assay Kit). Isolated cytoplasmic protein was incubated with acetyl-Asp-Glu-Val-Asp-7-amido-4-methyl coumarin or Ac-DEVD-AMC (50 μM final concentration) for 1 hour at room temperature in the dark. Caspase-3-like activity was measured by determining the cleavage of the fluorogenic caspase-3 substrate Ac-DEVD-AMC. Release of fluorescent AMC was measured with a 360 nm excitation wavelength and 460 nm emission wavelength in a TECAN Pro200 plate reader. Known amounts of hydrolyzed AMC was used to generate a calibration curve following the manufacturer’s instruction. Caspase enzymatic activity was calculated by finding the slope of the product’s concentration plotted as a function of time. This activity was then standardized to the quantity of total proteins present in the sample (Bradford assay, Bio-Rad). For inhibitor assays, samples were resuspended in the same assay buffer and preincubated for 1 h with a caspase inhibitor (Ac-DEVD-CHO; 100 μM) before subjecting the samples to the same enzymatic assay described above.

### Vacuolar pH

Vacuolar pH was measured as described (Kwon et al., 2018) with minor modifications. 10 μm BCECF-AM and 0.02% (v/v) pluronic F-127 (Invitrogen) were added to seedlings during the last hour of their equilibration in liquid MS medium containing 140 mM mannitol and 600 nM ISX, then incubated in the dark for 1 hour with gentle shaking. Before imposing stress treatments, seedlings were washed twice with same media but without the dye. The stained seedlings were then subjected to iso-osmotic or hypo-osmotic treatment as described above for 3 hours. Florescent images of stained roots were obtained using a confocal microscope (Olympus Fluoview FV3000) with excitation at 445 and 488 nm and emission at 525–550 nm. Fluorescence intensity in the elongation zone of roots obtained at 488 nm excitation was divided by that obtained at 445 nm excitation using Fiji (Schindelin et al., 2012). An *in situ* calibration was performed separately using wild type seedlings to make calibration curve as described by (Kwon et al., 2018). Relative change in vacuolar pH was calculated using the calibration curve.

### Monitoring ROS

Accumulation of ROS was assayed by confocal microscopy of seedlings stained with the fluorescent dye 2’,7’-dichlorofluorescin diacetate (H_2_DCFDA; Sigma-Aldrich) as described previously (Achard et al., 2008). Five-day-old seedlings were collected and incubated with 50 μM H_2_DCFDA dissolved in either MS supplemented with 140 mm mannitol or MS for 30 min, then washed three times with the appropriate medium. For the quantification of fluorescence H_2_DCFDA stained seedlings were excited at 488 nm; emitted light was detected at 505 to 530 nm. The green fluorescence in the elongation zone of roots was quantified using Fiji software (Schindelin et al., 2012). Fluorescent pixel intensity quantified after subtracting the pixel intensity of the background. Data are reported as relative fold change (fluorescence intensity of treated root)/ (fluorescence intensity of control root), to account for mechanical perturbations that may have occurred during the assay. For ROS inhibition, seedlings were incubated with the ROS inhibitors or scavengers, including KI (10 μM), DPI (10 μM) and SHAM (50 μM) as previously described (Chen et al., 1996; Rouet et al., 2006; Kurusu et al., 2012a). These chemicals were added during the 4-hour equilibration step before subjecting the seedlings to either iso or hypo-osmotic treatments.

### Measuring redox status using roGFP biosensors

For ratiometric analysis of roGFP1, seedlings expressing cytoplasmic roGFP1 were excited with 405 and 488 nm lasers, and emission was collected at 510 nm using a confocal laser scanning microscope (Olympus Fluoview FV 3000). Fluorescence intensity was determined from the transition/elongation zone of the seedlings subjected to cell swelling assay. The fluorescence ratios were obtained by dividing the intensities obtained at 405 nm by 488 nm using Fiji software (Schindelin et al., 2012). To calibrate roGFP1 probe, seedlings harboring roGFP1 with indicated genotypic backgrounds were treated with 10 mM DTT (full reduction) and with 10 mM H_2_O_2_ (complete oxidation), respectively as previously described (Jiang et al., 2006).

### Aequorin-based [Ca^2+^]_cyt_ luminescence

Three-day-old seedlings of wild type, *msl10-1* and *msl10-3G* mutant lines expressing *AEQ* were transferred individually to wells of sterile, white, 96-well microplates (Greiner Bio-One) containing 130 μL of 1x MS supplemented with 140 mM mannitol, then incubated in 2 μM native coelenterazine (Sigma) overnight, in darkness and at room temperature. The following day, 600 nM ISX (1.3 μl) was added to each well and the plate gently shaken for 4 hours. In a plate reader, the baseline luminescence was measured for 5 s, then either 150 μL of water or 140 mm mannitol were injected to produce hypo-osmotic or iso-osmotic treatments, respectively. The injection speed was set at 150 μL^s−1^. The luminescence emitted from the seedlings was measured for 30 s over a 1.1-s interval. Conversion of luminescent values to [Ca^2+^]_cyt_ was performed as described by (Knight et al., 1997). All results shown include data from four biological replicates. For each biological replicate, seven seedlings per transgenic lines were used. Three independent transgenic lines expressing aequorin in wild type, *msl10-1* and *msl10-3G* lines were used. In some cases, the *msl9-1* mutant allele was segregating in the *msl10-1* background, but we did not observe any difference in calcium response between siblings.

### Gene expression analysis

RNA was extracted using the RNeasy Plant Mini Kit with in-column DNase I digestion (Qiagen). Five hundred ng total RNA was reverse transcribed using oligo(dT) and the MLV reverse transcription system following the manufacturer’s instructions (Promega). qRT-PCR was performed in 25 μL reaction volumes with gene-specific primers and conducted with a StepOne Plus Real-time PCR System (Applied Biosystems) using the SYBR Green qPCR Master Mix (Thermofisher). Elongation factor 1α (*EF1*) and ubiquitin 10 (*UBQ10*) were used as reference genes for normalization of gene expression levels as previously described (Basu et al., 2020). Reported expression levels are expressed normalized to the wild type at 0 h or control treatment as indicated in the figure legends. Primer sequences are listed in **Supplemental Table 1**.

### Immunoblot analysis

Total protein extraction and immunoblotting was performed according to a method described previously (Veley et al., 2014; Basu et al., 2020). 12- to 18-day-old seedlings were collected in a 2 mL microcentrifuge tube, weighed, frozen in liquid nitrogen, homogenized with pestle in 2X sample buffer, and denatured for 10 min at 70°C. Proteins samples were resolved by 10% SDS-PAGE and transferred to polyvinylidene difluoride membranes (Millipore) for 16 h at 100 mA. Transferred proteins were first probed with anti-GFP (Takara Bio, 1:5,000 dilution) for 16 h. Next, membrane was reprobed with anti-tubulin (Sigma, 1:20,000 dilution) antibodies with incubation for 2 h. In both cases, 1 h incubation with horseradish peroxidase-conjugated anti-mouse secondary antibodies (1:10,000 dilution; Millipore) was performed. Detection was performed using the SuperSignal West Dura Detection Kit (Thermo Fisher Scientific).

### Statistical analyses

Graphing and statistical analysis was performed with GraphPad Prism version 7.04. In all cases, significant differences refer to statistical significance at P ≤ 0.05. The specific statistics applied in each experiment are indicated in the corresponding figure legends.

### Accession Numbers

The accession numbers for the genes discussed in this article are as follows:

*MSL10 (At5G12080), PDF1.2 (At5G44420), PERX34 (At3g49120), UBQ5 (At3g62250), EF1α (At5g60390), MC1 (AT1G02170), MC2 (AT4G25110), MC9 (AT5G04200), WRKY18 (AT4G31800), TCH1 (AT5G37780), TCH2 (AT5G37770), TCH3 (AT2G41100), TCH4*, BFN1 (*AT1G11190), RNS3 (AT1G26820), TCH4 (AT5G57560)*.

## AUTHOR CONTRIBUTIONS

D.B. and E.S.H. conceived the project and designed the experiments; D.B. performed the experiments; E.S.H created the figures; D.B. and E.S.H. wrote the manuscript.

## ACKNOWLEDGMENTS

We thank the Arabidopsis Biological Resource Center for pBINU-CYA. This work was supported by the NSF Science and Technology Center Grant 1548571, NASA NNX13AM55G to E.S.H, and NSF MCB 1253103 to E.S.H. No conflicts of interest declared.

---

**Figure S1.**
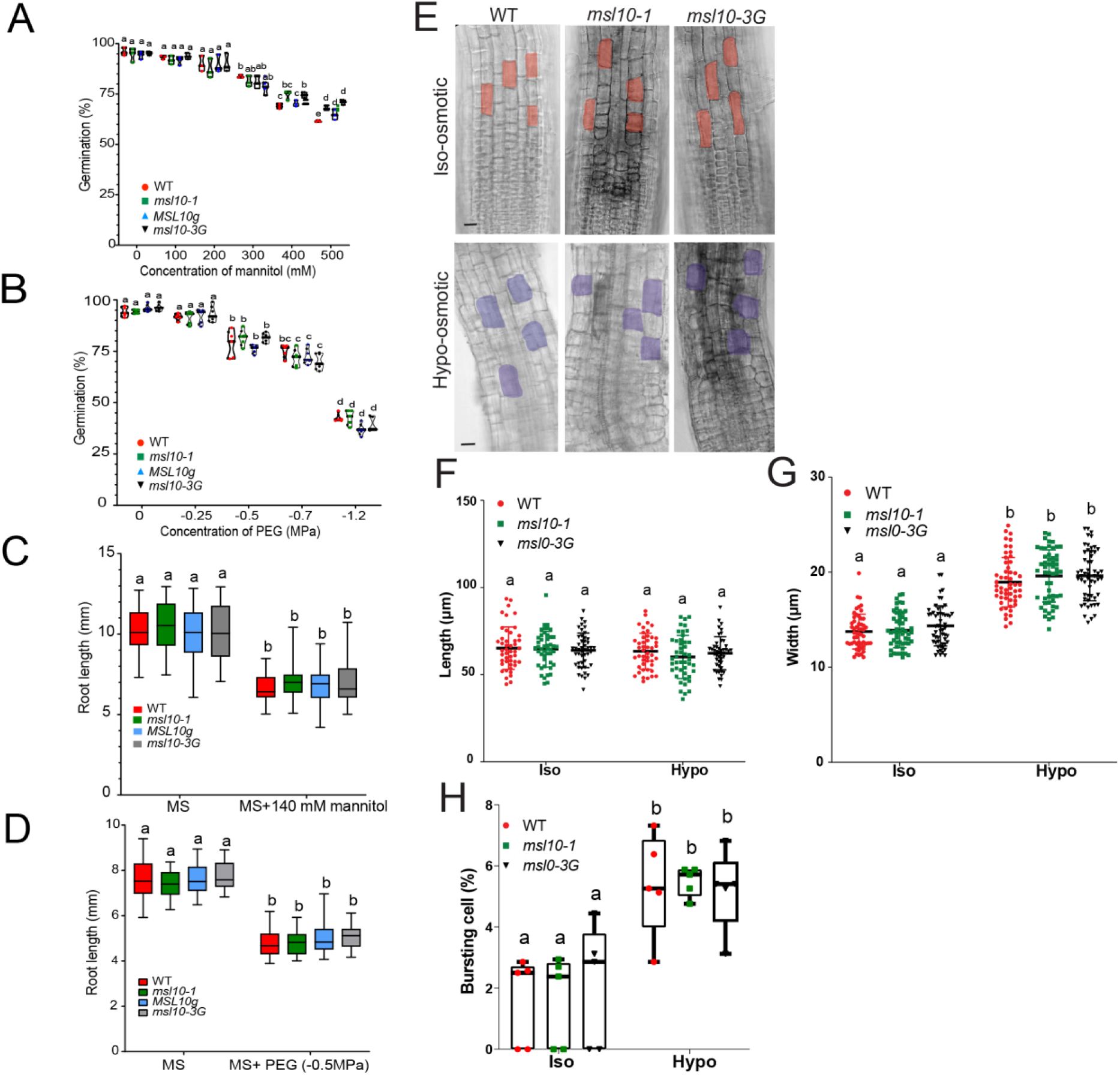
Establishing a seedling-based cell swelling assay to probe the function of MSL10. **(A)** and **(B)** Seed germination on increasing mannitol (mM) and PEG (MPa) treatments. ~150 seeds for each line were sterilized and sown on plates containing 1X MS medium with or without the indicated concentrations of mannitol or 1/2X MS with or without the indicated concentrations of PEG-8000 (MPa) as described (van der Weele et al., 2000). Seeds with protruded radicles were deemed germinated. **(C)** and **(D)** Primary root growth on140 mM mannitol or −0.5MPa PEG compared to IX-strength MS media. ~15 to 20 seeds for each line were sown and grown vertically for 7 days. Fifteen to nineteen seedlings per line were measured. Plates were scanned using the Syngene PXi imagining system equipped with GeneSys image acquisition software, and root lengths were measured using ImageJ. **(E)** Representative single optical slice of confocal images of five-day-old seedlings. Seedlings were subjected to osmotic down shock as described in Figure 1A and mounted and imaged within 30 seconds. Colored outlines indicate cells from root transition/elongation zone that were used for quantification in **(F)** and **(G)**. Scale bars, 10 μm. **(F), (G)** and **(H)** Scatter plot showing cell length, width and bursting (%) in the root transition zone of seedlings of indicated genotypes (n=9-11 cells per seedling; total of 54 cells per genotype). Seedlings were treated as described in (E). The length and width were measured using ImageJ. Data are means and SD from two independent experiments. **(A), (B), (C), (D), (F)** and **(G)** Two-way ANOVA with posthoc Tukey’s test was used to assess statistical differences; different letters denote significant differences (P < D.05). Data are means and SD from three independent experiments.

**Figure S2.**
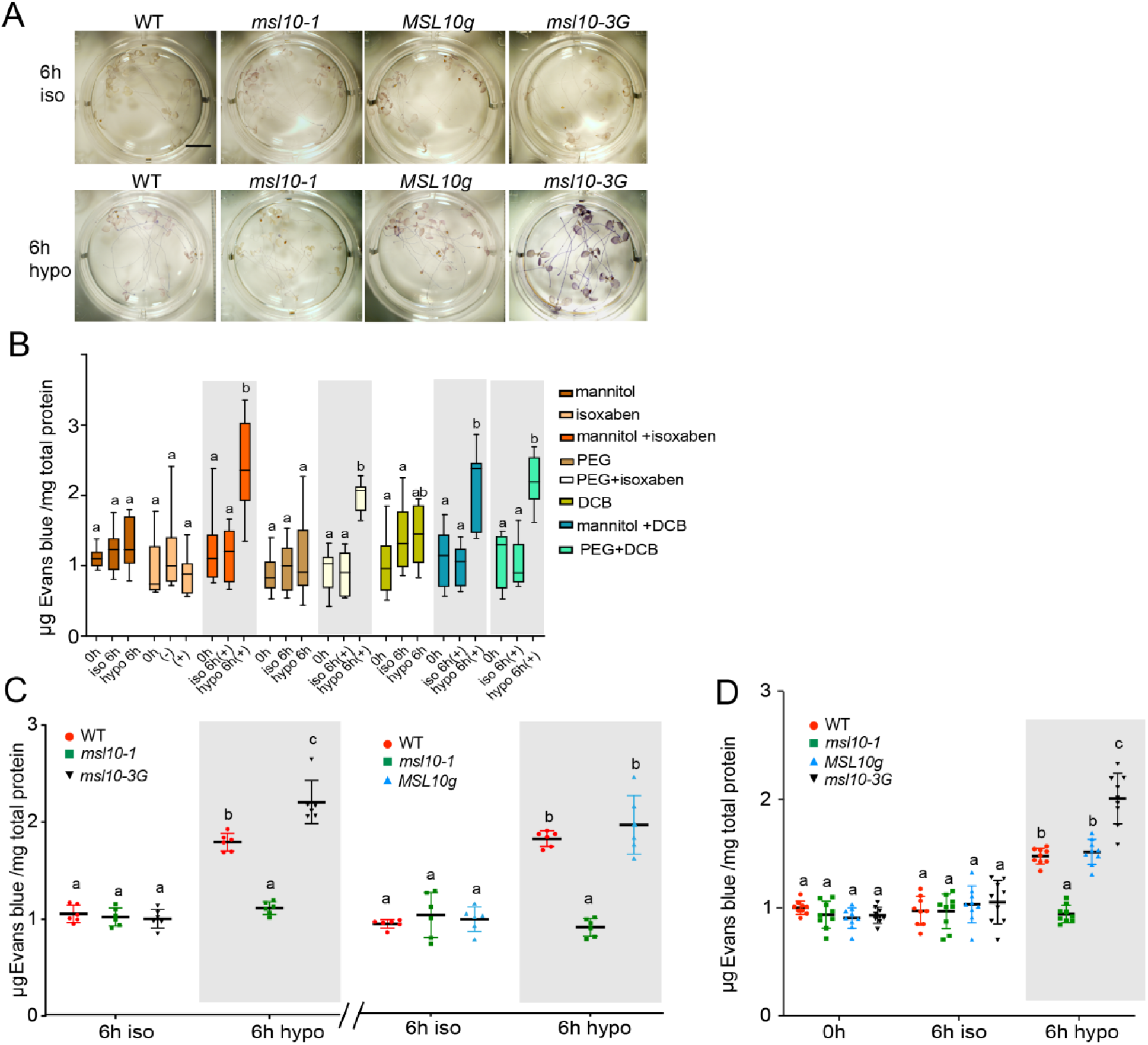
Monitoring MSL10-associated cell death in response to cell swelling. **(A)** Representative image of Evans blue-stained seedlings subjected to iso- or hypo-osmotic stress as in Figure 1 A. Seedlings were stained with D.25% (w/v) Evans blue for 15 min with gentle shaking. Seedlings were destained in an ascending series of aqueous ethanol solutions and resuspended in 10% glycerol. Images were captured with an Olympus DP8O equipped with a cooled color digital camera. **(B)** Cell death in wild type seedlings in response to varying treatments. Five-day-old seedlings were subjected to osmotic down shock after growth on 1X MS + PEG −(0.5MPa) or 1X MS + mannitol (140 mM) to 1XMS, disruption of cellulose biosynthesis by incubating with ISX (600 ⊓M) or DCB (500 nM), or by ∞mbining osmotic down shock with cell wall disruption treatments. Cell death was estimated spectroscopically by measuring the absorbance at 630 nm and normalized to total protein at 595 nm. **(C)** Cell death in response to a large osmotic downshock only, as assessed by Evans blue staining. Six-day-old seedlings of indicated genotypes were treated with osmotic down shock by transferring seedlings from 1X MS supplemented with 330 mM mannitol to 1X strength MS media. **(D)** Effect of shaking on swelling-induced cell death as assessed by Evans blue staining. Six-day-old seedlings of indicated genotypes were grown on 140 mM mannitol and subjected to the cell swelling assay as described in Figure 1 and the methods, but without shaking. **(A), (B), (C)**, and **(D)** Two-way ANOVA with posthoc Tukey’s test was used to assess statistical differences; different letters denote significant differences (P < 0.05). Data are means and SD from three independent experiments. Scale bar=1 mm.

**Figure S3.**
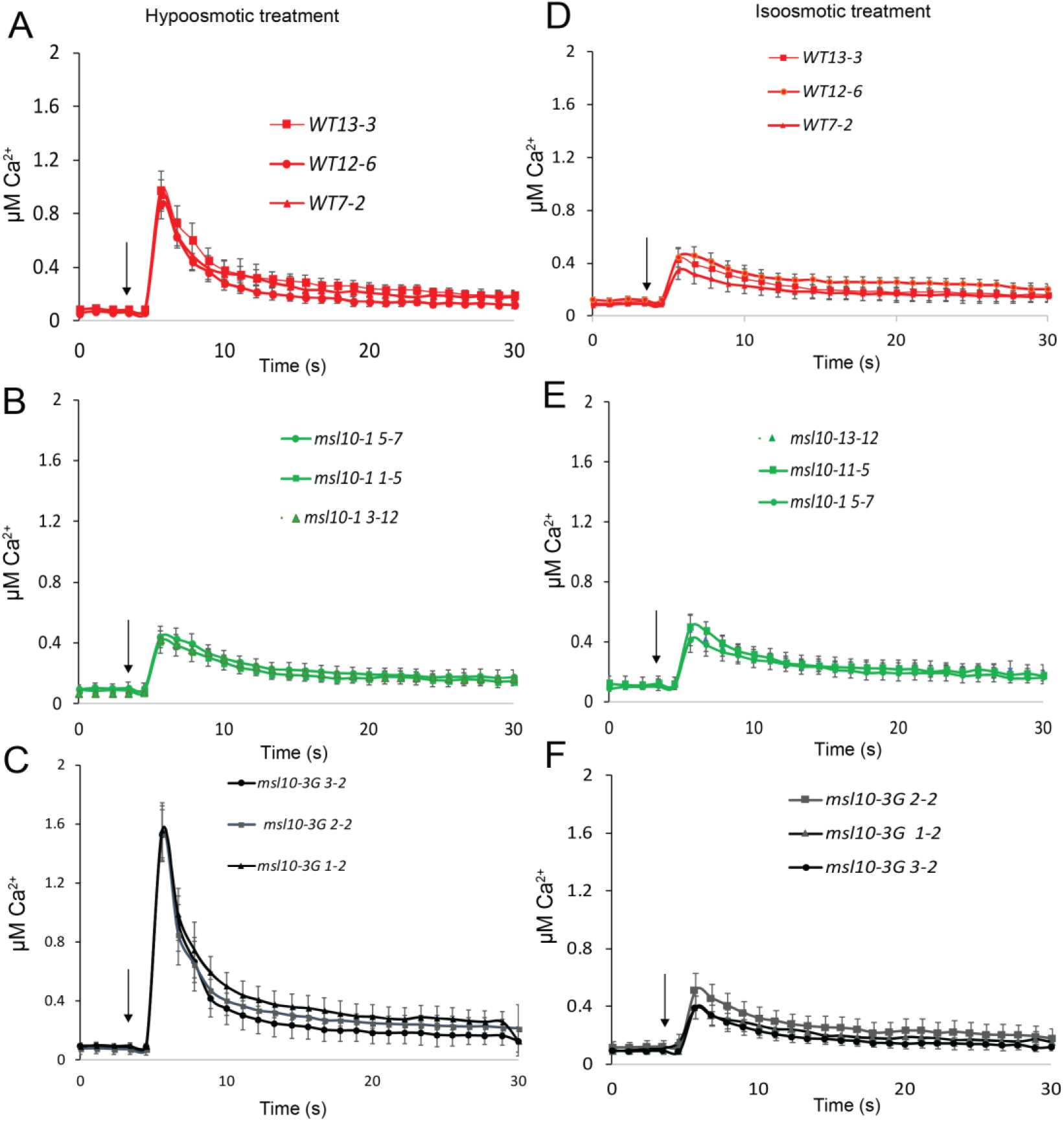
MSL10-dependent increases in swelling-induced intracellular Ca^2+^transients. Traces showing elevated Ca^2+^transient in seedlings harboring reconstituted aequorin after hypo-osmotic **(A-C)** or iso-osmotic **(D-F)** shock. Three independent T3 homozygous transgenic lines per genetic backround expressing aequorin were used in this experiment. Three- or four-day-old transgenic seedlings were grown on solid MS plates supplemented with 140 mM mannitol and aequorin reconstituted overnight with 2 μM coelenterazine in the dark. Seedlings were incubated with ISX (600 nM) for 4h with gentle shaking in the dark. Then, intracellular Ca^2+^ response was monitored by luminometry using a plate reader and either deionized water (hypoosmotic condition) or 140 mM mannitol (isosmotic condition) injected into the media (arrows). Values plotted in the traces correspond to the means of four independent experiments, seven seedlings per transgenic line, per treatment. Error bar represents ± SEs.

**Figure S4.**
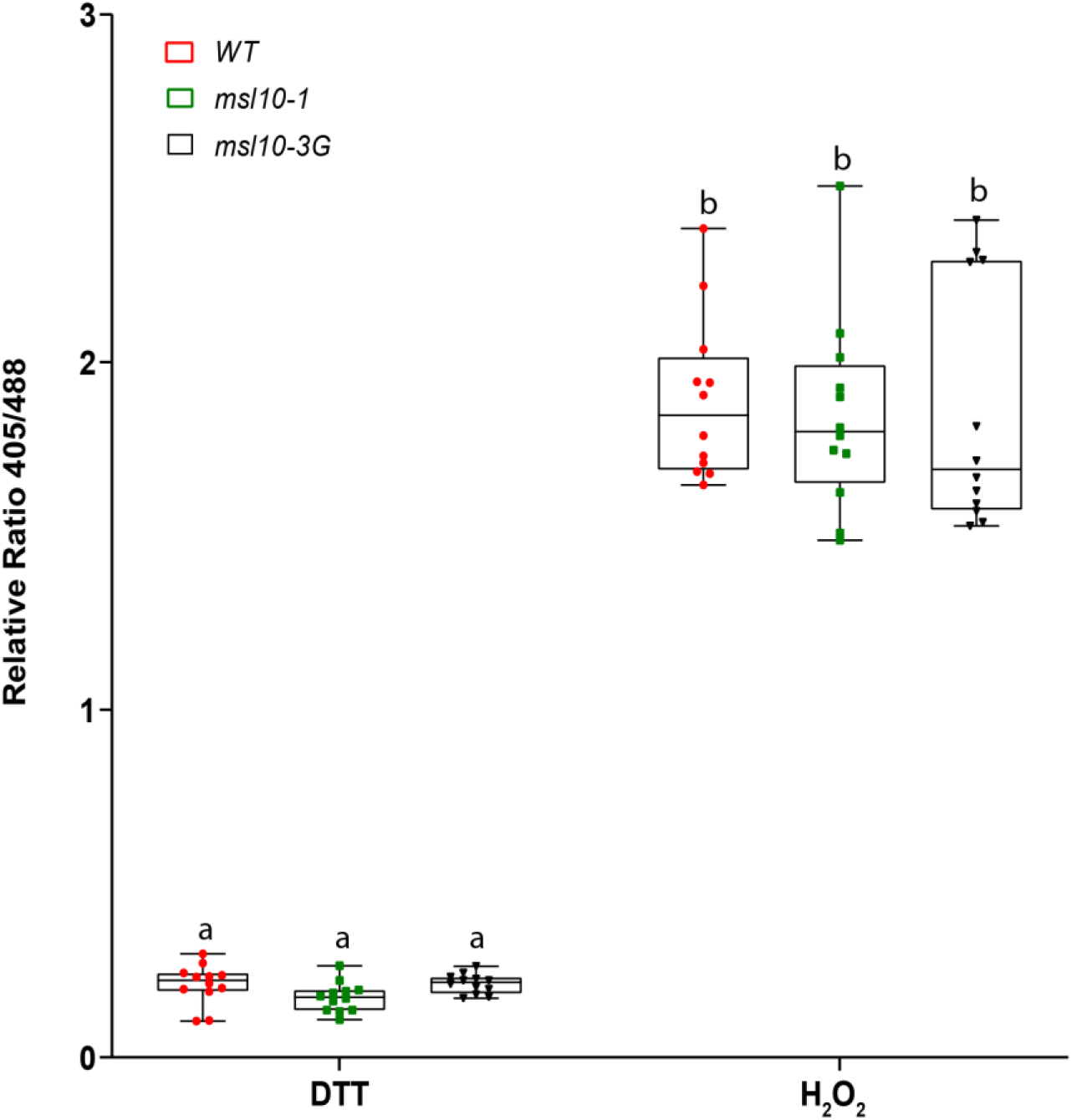
Determining the full dynamic range of redox-sensitive roGFP from seedlings expressing *roGFP1*. To calibrate the roGFP1 probe, seedlings harboring *roGFP1* were treated with 10 mM DTT (full reduction) or with 10 mM H_2_O_2_ (complete oxidation).The ratio of emmission at 510 nm after excitation at 405 nm or 488 nm values was determined from 12 individual seedlings for each condition. Error bars represent SD.Two-way ANOVA with posthoc Tukey’s test was used to assess statistical differences; identical letters denote no significant differences (P < 0.05).

**Figure S5.**
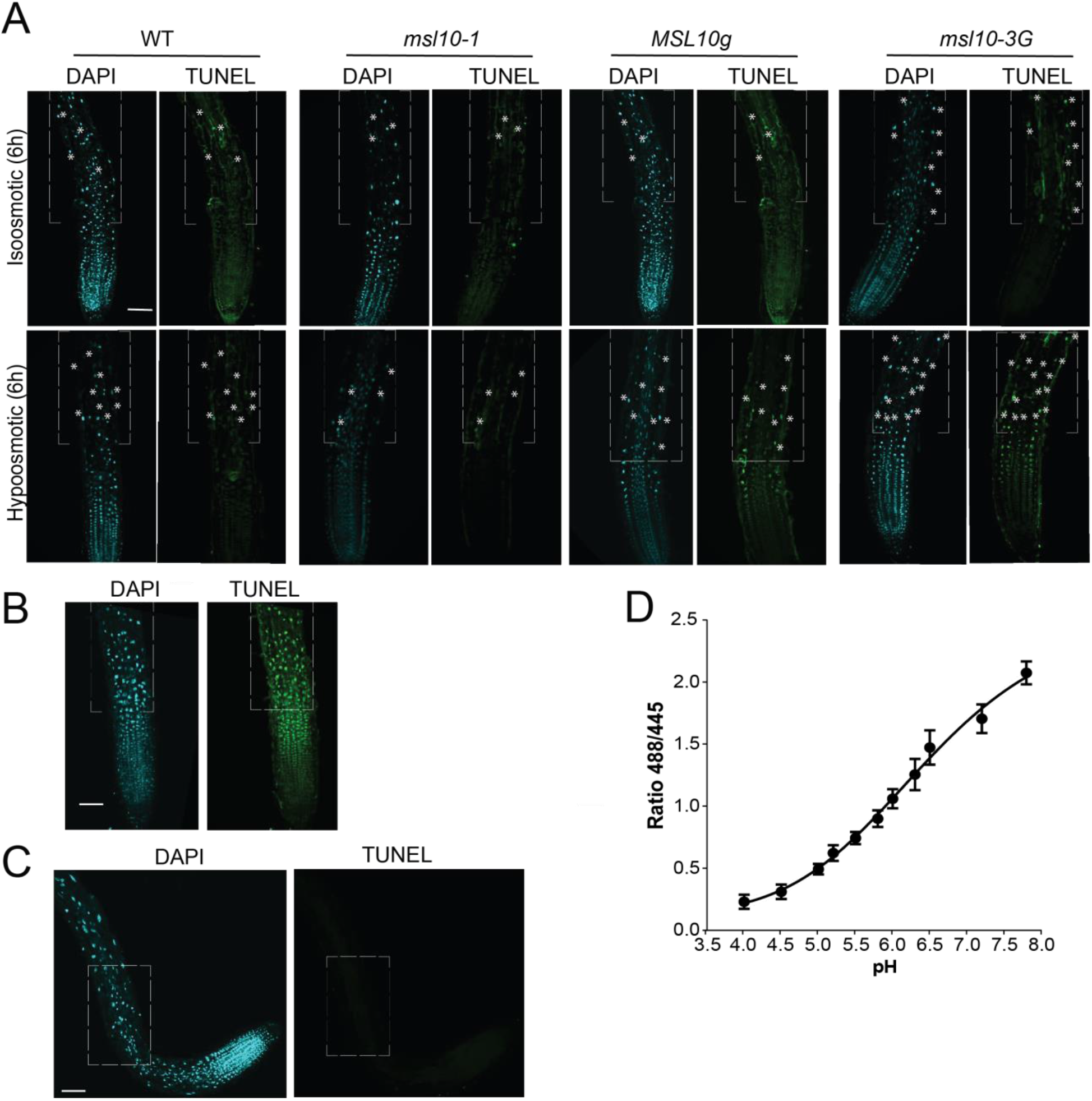
Representative images of TUNEL-stained seedling roots and pH calibration curve of ratiometric pH indicator BCECF-AM. **(A)** Roots of seedlings after TUNEL and DAPI staining. Five-day-old seedlings of the indicated genotypes were subjected to the swelling assay as in Figure 1A. TUNEL-positive nuclei (in green) and DAPI stained (in blue) nuclei are shown. Asterisks indicate TUNEL-positive nuclei. Boxes indicate the area of the root elongation zone used for counting TUNEL-positive nuclei as shown in Figure 5. **(B)** Typical fluorescent image of primary root treated with DNase I for 15 min before TUNEL staining. **(C)** Typical fluorescent image of primary root incubated without terminal transferase enzyme. **(A), (B)** and **(C)** Images were captured by a confocal laser scanning microscope (Olympus Fluoview FV 3000). Scale bar = 50 μm. **(D)** *In situ* calibration curve of the pH of BCECF-AM dye measured from elongation zone of root.To generate a pH calibration curve, five-day-old seedlings were incubated for 20 min in pH equilibration buffer containing either 50 mm MES-BTP (pH 4-6.5) or 50 mm HEPES-BTP (pH 7-7.8) and 50 mm ammonium acetate as described (Zhu et al., 2018). Fluorescent images of stained roots were obtained using a confocal microscope (Olympus Fluoview FV 3000) after excitation with 445 and 488 nm and emission collected at 525-550 nm. The intensity obtained at 488 nm excitation was divided by that obtained at 445 nm excitation.The average ratio values were determined from 12 individual seedlings for each condition.

**Figure S6.**
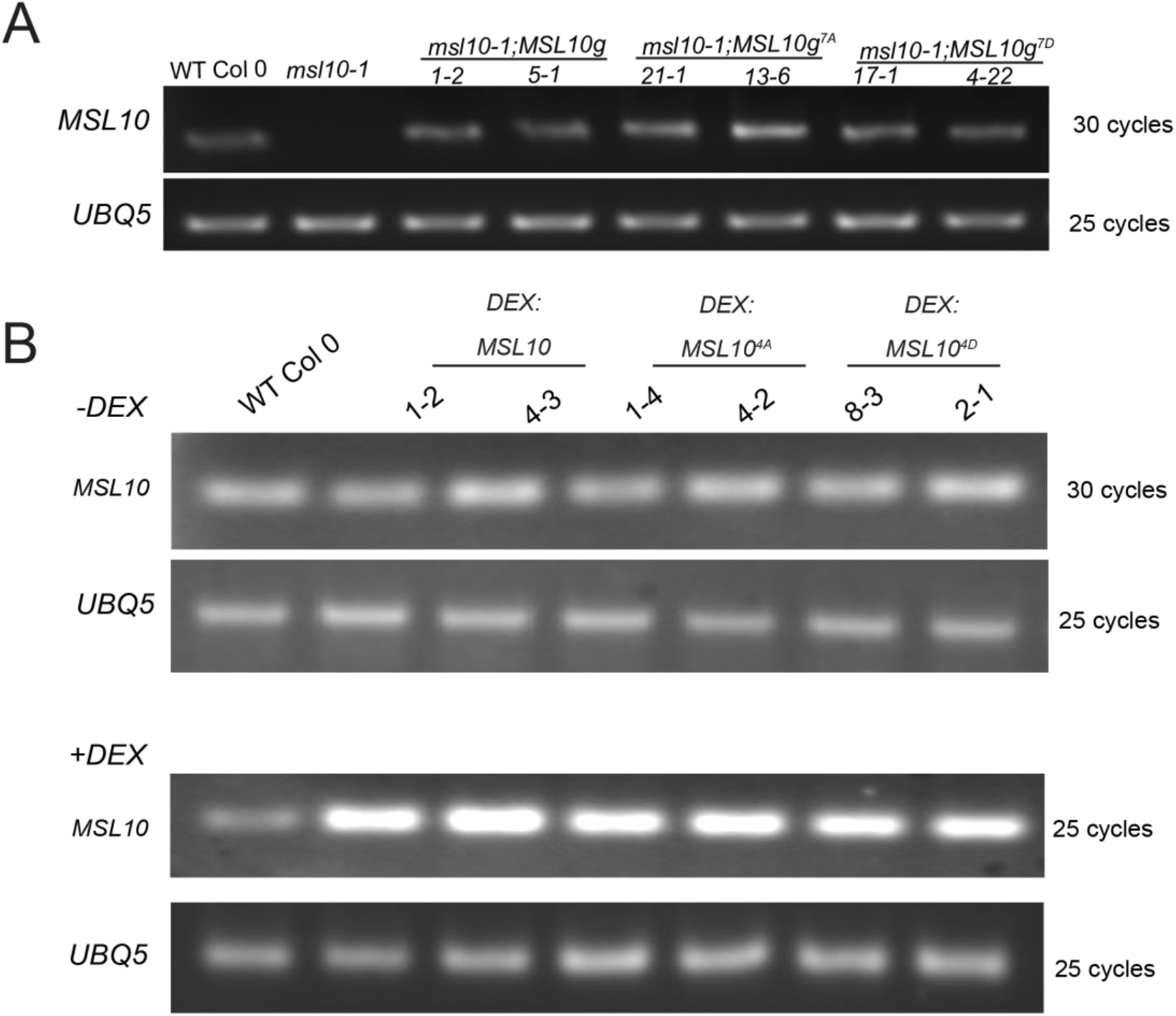
Monitoring *MSL10* transcript levels in seedlings grown on 1X MS supplemented with 140 mM mannitol using semi-quantitative RT-PCR. **(A)** Expression profiles of *MSL10* from *msl10-1* lines harboring genomic version of *MSL10* and its phospho variants. (B) Expression profiles of *MSL10* before and after DEX treatment of transgenic lines expressing *DEX: MSL10* and its phospho-variants. Total RNA was isolated from eighteen five-day-old seedlings of wild type, *msl10-1*, and two independent homozygous T3 lines either expressing genomic version of *MSL10* or harboring DEX inducible *MSL10* and its phospho variants. The expression of *UBQ5* was used as a control. For conditional induction of *MSL10*, seedlings were incubated with 30 μM DEX for 12 h.

**Supplemental Table 1.**
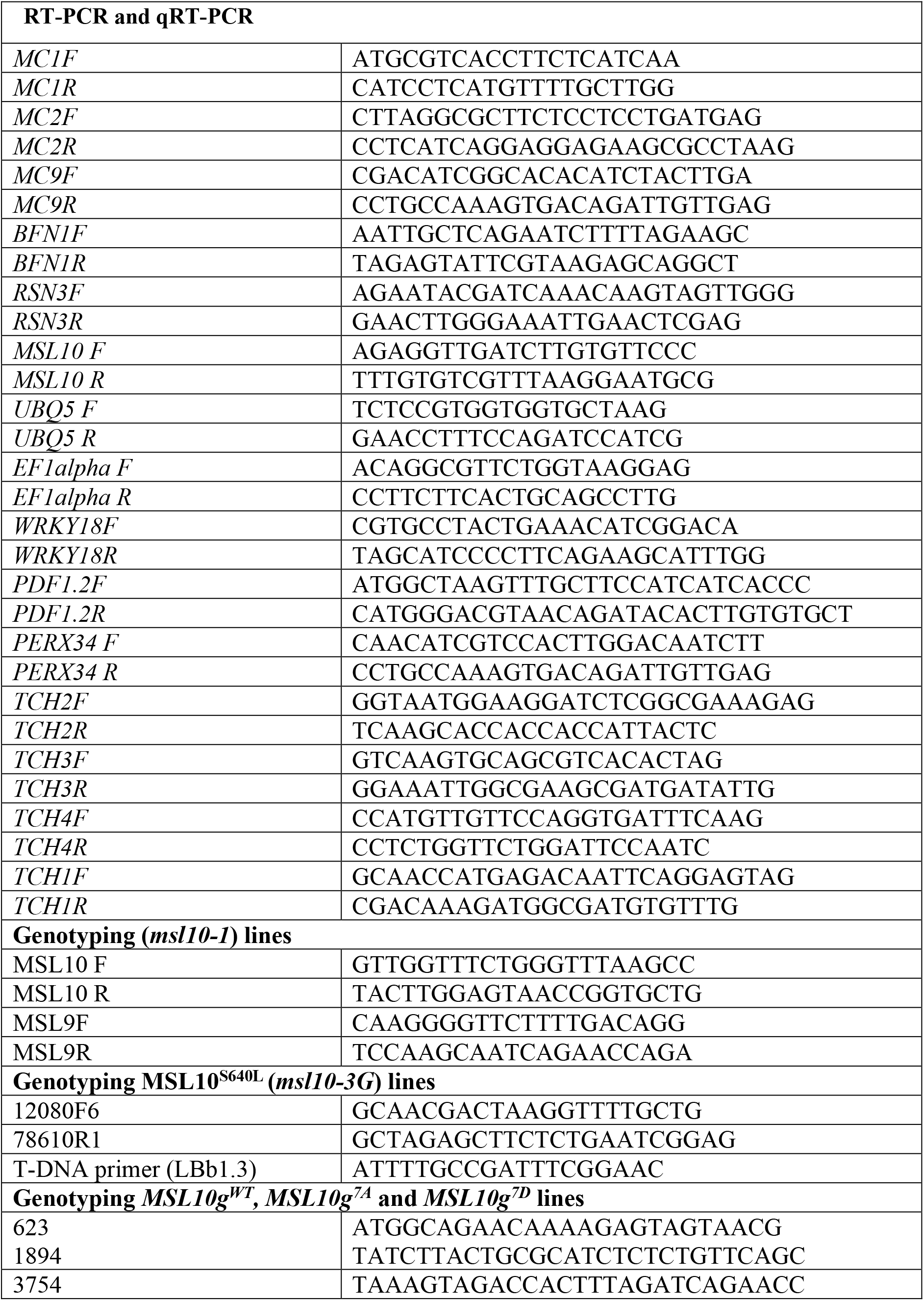
List of primers used in this study.

